# Epigenetic Programming of Estrogen Receptor in Adipose Tissue by High Fat Diet Regulates Obesity-Induced Inflammation

**DOI:** 10.1101/2025.06.21.660886

**Authors:** Rui Wu, Fenfen Li, Shirong Wang, Jia Jing, Xin Cui, Yifei Huang, Xucheng Zhang, Jose A. Carrillo, Zufeng Ding, Jiuzhou Song, Liqing Yu, Huidong Shi, Bingzhong Xue, Hang Shi

## Abstract

Adipose inflammation plays a key role in obesity-induced metabolic abnormalities. Epigenetic regulation, including DNA methylation, is a molecular link between environmental factors and complex diseases. Here we found that high fat diet (HFD) feeding induced a dynamic change of DNA methylome in mouse white adipose tissue (WAT) analyzed by reduced representative bisulfite sequencing. Interestingly, DNA methylation at the promoter of estrogen receptor α (*Esr1*) was significantly increased by HFD, concomitant with a down-regulation of *Esr1* expression. HFD feeding in mice increased the expression of DNA methyltransferase 1 (*Dnmt1*) and *Dnmt3a,* and binding of DNMT1 and DNMT3a to *Esr1* promoter in WAT. Mice with adipocyte-specific *Dnmt1* deficiency displayed increased *Esr1* expression, decreased adipose inflammation and improved insulin sensitivity upon HFD challenge; while mice with adipocyte-specific *Dnmt3a* deficiency showed a mild metabolic phenotype. Using a modified CRISPR/RNA-guided system to specifically target DNA methylation at the *Esr1* promoter in WAT, we found that reducing DNA methylation at *Esr1* promoter increased *Esr1* expression, decreased adipose inflammation and improved insulin sensitivity in HFD-challenged mice. Our study demonstrates that DNA methylation at *Esr1* promoter plays an important role in regulating adipose inflammation, which may contribute to obesity-induced insulin resistance.

## INTRODUCTION

Obesity is characterized by chronic inflammation that causally links obesity to insulin resistance/type 2 diabetes (1, 2). Adipose tissue plays a key role in the generation of inflammatory response and mediators in obesity (1, 2). Inflammatory signaling pathways in fat tissue can be activated in nutrient-rich conditions (3). However, it is still not fully understood how inflammatory program is altered by nutrient-rich conditions that often promotes obesity.

Since the original reports linking chronic inflammation in adipose tissue to obesity-induced insulin resistance, there have been numerous publications studying pathways underlying obesity-induced inflammation in metabolic dysfunctions (2). While most studies have been devoted to the evaluation of genetic pathways (eg ER stress, oxidative stress, ceramide formation) that mediates nutrient-induced abnormalities in metabolic tissues (2), much less is known about epigenetic mechanisms, a link between environmental factors (e.g. diets) and complex diseases (e.g. obesity and diabetes), in this process. One of the most common epigenetic regulations is DNA methylation of cytosines at mainly CpG dinucleotides, which frequently takes place in the promoter and 5’ regions of genes (4). DNA hypomethylation at gene promoters often results in transcriptional activation, while hypermethylation is often associated with gene silencing (4). *De novo* DNA methylation is mediated by DNA methyltransferase (DNMT) 3a and 3b. Once established, DNA methylation is then maintained through mitosis primarily by the maintenance enzyme DNMT1 (5, 6). However, recent evidence also supported a role of DNMT1 in *de novo* DNA methylation (7). On the other hand, DNA demethylation can be achieved by the ten-eleven translocation 1 (TET1) dioxygenase that catalyzes the hydroxylation of 5-methylcytosine (5-mC) to 5-hydroxymethylcytosine (5-hmC) and subsequent generation of 5-formylcytosine (5-fC) and 5-carboxylcytosine (5-caC), which are then converted to unmodified cysteines by replication-related dilution or glycosylation-mediated base-excision repair (8, 9). The key role of DNA methylation and its maintenance enzyme DNMT1 in metabolic inflammation has been revealed in our recent report demonstrating that hypermethylation at peroxisome proliferator activated receptor γ (*Pparγ)* promoter by saturated fatty acids and proinflammatory cytokines, levels of which are commonly elevated in obesity, promotes M1 proinflammatory macrophage polarization and inflammation in adipose tissue, resulting in insulin resistance in obesity (10). However, it is not clear whether and how DNA methylation plays a role in adipocyte chemotaxis and inflammation, an integral part of overall adipose inflammation.

In the present study, we tested the hypothesis that DNA methylation at estrogen receptor α (*Esr1)* promoter mediates adipocyte inflammation and chemotaxis, leading to obesity-induced insulin resistance. Using reduced representation bisulfite sequencing (RRBS) analysis, we conducted a genome-wide profiling of DNA methylation in adipose tissue of diet-induced obese (DIO) mice. From the RRBS analysis, we discovered a significant increase in DNA methylation at *Esr1* promoter, which is associated with down-regulation of *Esr1* expression. We also found that *Esr1* promoter activity was differentially regulated in unmethylated vs. methylated state. Using ChIP assays, we found an increased binding of DNMT1 and DNMT3a to the *Esr1* promoter in white adipose tissue of mice fed high fat diet (HFD). We then generated adipocyte-specific *Dnnmt1* and *3a* knockout (AD1KO and AD3aKO) mice and characterized their inflammatory status and metabolic phenotypes. Finally, we used a novel CRISPR/RNA-guided system to specifically induce methylation/demethylation at the *Esr1* promoter and further characterized adipocyte inflammation/chemotaxis, macrophage infiltration and insulin sensitivity in these mice challenged with HFD feeding.

## RESULTS

### Dynamic changes of the DNA methylome in white fat of HFD-fed mice

To study whether high fat diet (HFD) reprograms DNA methylome in fat tissue, we performed a DNA methylation profiling experiment in gonadal white fat (gWAT) of male mice fed with either a low fat diet (LFD) or HFD for 12 weeks using Reduced Representation Bisulfite Sequencing (RRBS) approach (11-13). Our bioinformatic analysis showed that there were up to 1630 Differentially Methylated Regions (DMRs) in HFD-vs chow-fed mice. Around 44.1% of the DMRs were located in intergenic regions, with the rest of the DMRs distributed within genes including 5’-end/5’-untranslated region (5’-UTR), cDNA coding sequences (CDS), introns and 3’-end/3’-UTR regions (**Figure 1A**). Within the genes with altered DNA methylation level, 700 genes exhibited up-regulated DNA methylation rates by HFD and 211 genes exhibited down-regulated DNA methylation rates by HFD. These data suggest that HFD feeding dynamically changes DNA methylome in fat, with the majority of the genes (77%) exhibit an increase in DNA methylation status. In addition, Gene Ontology (GO) and Kyoto Encyclopedia of Genes and Genomes (KEGG) analysis suggested that genes with methylation changes were involved in various pathways including biological process, molecular function and cellular component (**Supp. Fig 1**).

**Figure 1.**
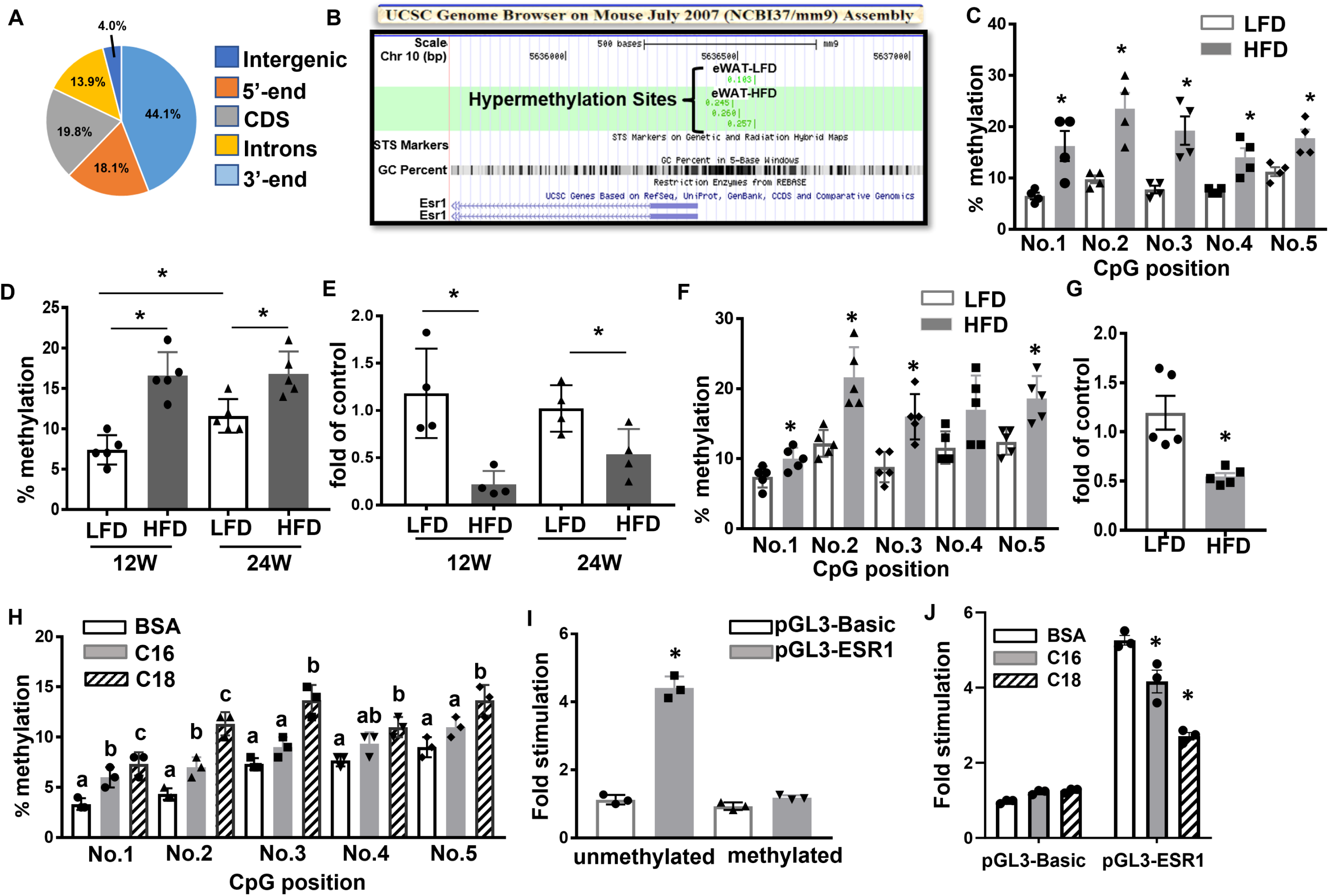
High fat diet (HFD) regulates *Esr1* expression via promoter DNA methylation. (A) Distribution of DMRs in the genome in gWAT of male mice fed with LFD or HFD for 12 weeks. (B) RRBS profiling of DNA methylation levels at *Esr1* promoter in gWAT of LFD- and HFD-fed mice. RRBS was performed in gWAT from male C57BL/6J mice fed with either LFD or HFD for 12 weeks starting at 6 weeks of age. Genomic DNA from 4 animals was pooled in each group for RRBS analysis. (C) DNA methylation levels at individual CpG sites at *Esr1* promoter in gWAT of male C57BL/6J mice fed either LFD or HFD for 12 weeks, n=4/group. *p<0.05 vs. LFD by Student’s t test. (D-E) Average DNA methylation levels at *Esr1* promoter (D, n=5/group) and Esr1 expression (E, n=4/group) in gWAT of male C57BL/6J mice fed either LFD or HFD for 12 or 24 weeks. *p<0.05 vs. LFD by Student’s t test. (F-G) DNA methylation levels at individual CpG sites at *Esr1* promoter (F) and *Esr1* expression (G) in gWAT of female C57BL/6J mice fed either LFD or HFD for 12 weeks, n=5/group. *p<0.05 vs. LFD by Student’s t test. (H) DNA methylation levels at individual CpG sites at *Esr1* promoter in 3T3-L1 adipocytes treated with palmitate (C16) or stearate (C18), n=3/group. Groups labeled with different letters are statistically different from each other as analyzed by ANOVA with Fisher’s least significant difference (LSD) post hoc test. (I) *Esr1* promoter luciferase activity in 3T3-L1 adipocytes transfected with fully methylated or unmethylated constructs, n=3. *p<0.05 vs. all other groups as analyzed by ANOVA with Fisher’s LSD post hoc test. (J) *Esr1* promoter luciferase activity in 3T3-L1 adipocytes transfected with unmethylated constructs in the presence of BSA, palmitate (C:16, 200μM), or stearate (C:18, 200μM). n=3. *p<0.05 vs. BSA by ANOVA with Fisher’s LSD post hoc test. All data are expressed as mean ± SEM.

### Methylation of the Esr1 promoter is enhanced by HFD feeding possibly via DNMT1 and DNMT3a

Notably, our RRBS profiling indicated that methylation rate at the 5’-end of the *Esr1* promoter was significantly increased in gWAT of HFD-fed mice as indicated using the University of California Santa Cruz (UCSC) Genome Browser (**Fig 1B**). Since adipocyte *Esr1* has emerged as a novel anti-inflammatory molecule in obesity (14-16), we focus on *Esr1* as the major methylation target during diet-induced obesity in this study. Interestingly, the proximal promoter and 5’-UTR regions of *Esr1* are enriched with CpG islands, among which 5 CpG sites are located downstream adjacent to the TATA box (**Suppl. Fig 2**), raising a possibility that the *Esr1* promoter might be subjected to epigenetic regulation through DNA methylation by HFD. Indeed, our pyrosequencing analysis indicated that HFD feeding significantly increased DNA methylation at several CpG sites at *Esr1* promoter in gWAT of male C57/BL6J mice (**Fig 1C**). In consistency, the average DNA methylation rate at *Esr1* promoter in gWAT was increased in both 12-week and 24-week HFD-fed mice (**Fig 1D**). This was associated with down-regulation of *Esr1* mRNA expression in gWAT of both 12-week and 24-week HFD-fed mice, with a more profound reduction observed in 12-week HFD-fed mice (**Fig 1E**). Interestingly, we observed that average DNA methylation level at *Esr1* promoter was already increased in 24-week LFD-fed mice compared to that of 12-week (**Fig 1D**), suggesting that besides dietary factors, aging may be another important factor that regulates DNA methylation at *Esr1* promoter, which may contribute to aging-associated metabolic dysfunction. This age-related change may contribute to the more profoundly increased *Esr1* promoter DNA methylation and more profoundly reduced *Esr1* expression observed between HFD- and LFD-fed mice at 12-weeks compared to that of 24-weeks (**Fig 1D-E**). Similarly, we also found that HFD-fed female mice, like male mice, also displayed increased DNA methylation rate at *Esr1* promoter coupled with decreased *Esr1* expression in the gonadal fat depots (gWAT) (**Fig 1F** and **1G**).

We then tested whether saturated free fatty acids (FFAs), whose levels are abundant in HFDs and are commonly elevated in obesity, regulates DNA methylation levels at the *Esr1* promoter. Primary preadipocytes were differentiated and treated with palmitate (C16, 200µM) and stearate (C18, 200µM) for two days. Pyrosequencing analysis showed that stearate significantly upregulated methylation rates at individual CpG sites at *Esr1* promoter, while palmitate exerted a lesser effect (**Fig 1H**). To determine whether *Esr1* promoter activity is indeed regulated by methylation, we cloned a 1kb proximal promoter regions at the *Esr1* locus including the CpG-enriched regions into pGL3-luciferase expression vector and examined the fully methylated vs unmethylated *Esr1* promoter activity by transfecting these constructs into 3T3-L1 cells. Our luciferase assays showed that the luciferase activity of the unmethylated promoter was more than 4-fold higher than that of the fully methylated promoter (**Fig 1I**). Interestingly, the increased luciferase activity in unmethylated *Esr1* promoter was significantly suppressed in the presence of 200μM palmitate or stearate (**Fig 1J**), further confirming that saturated fatty acids suppress *Esr1* promoter activity via regulating *Esr1* promoter DNA methylation.

To determine which DNA methyltransferases (DNMT1, 3A or 3B) and demethylases (TET1, 2 and 3) may mediate the increased DNA methylation at the *Esr1* promoter caused by HFD feeding, we examined the binding of DNMTs to the *Esr1* promoter using ChIP assay in gWAT of mice fed with either LFD or HFD. We found that HFD feeding significantly enhanced the binding of DNMT1 and DNMT3A to the *Esr1* promoter in gWAT (**Fig 2A-B**), while we could not detect any DNMT3B binding to *Esr1* promoter due to low expression (data not shown). Interestingly, HFD feeding also increased mRNA expression of *Dnmt1* and *Dnmt3a* in gWAT of HFD-fed mice (**Fig 2C-D**). A similar increase was also observed in DNMT1 protein levels in gWAT of HFD-fed mice (**Fig 2E**). Moreover, HFD feeding also down-regulated the expression of *Tet1* and *Tet2,* and tended to down-regulate *Tet3* expression in gWAT (**Suppl**. **Fig 3**). To further narrow down whether DNMT1 or 3A mediates *Esr1* promoter methylation, we knocked down *Dnmt1* or *Dnmt3a* in 3T3-L1 adipocytes after day 5 or day 8 of differentiation. We found that *Dnmt1* knockdown markedly promoted the expression of *Esr1* mRNA in adipocytes with *Dnmt1* knockdown at either day 5 or day 8 of differentiation, while *Dnmt3a* knockdown either had no effect on or even decreased *Esr1* expression (**Fig 2F-G**). To further study how *Dnmt1* regulates *Esr1* expression in more physiologically relevant settings, we generated adipocyte-specific *Dnmt1* knockout mice (AD1KO) by crossing *Dnmt1*-floxed mice (17) with adiponectin-Cre mice (18). We found that adipocytes differentiated from AD1KO mice significantly increased the expression of *Esr1* compared to adipocytes differentiated from fl/fl mice (**Fig 2H**). These data suggest that DNMT1 may be a key enzyme in mediating *Esr1* promoter methylation induced by HFD feeding.

**Figure 2.**
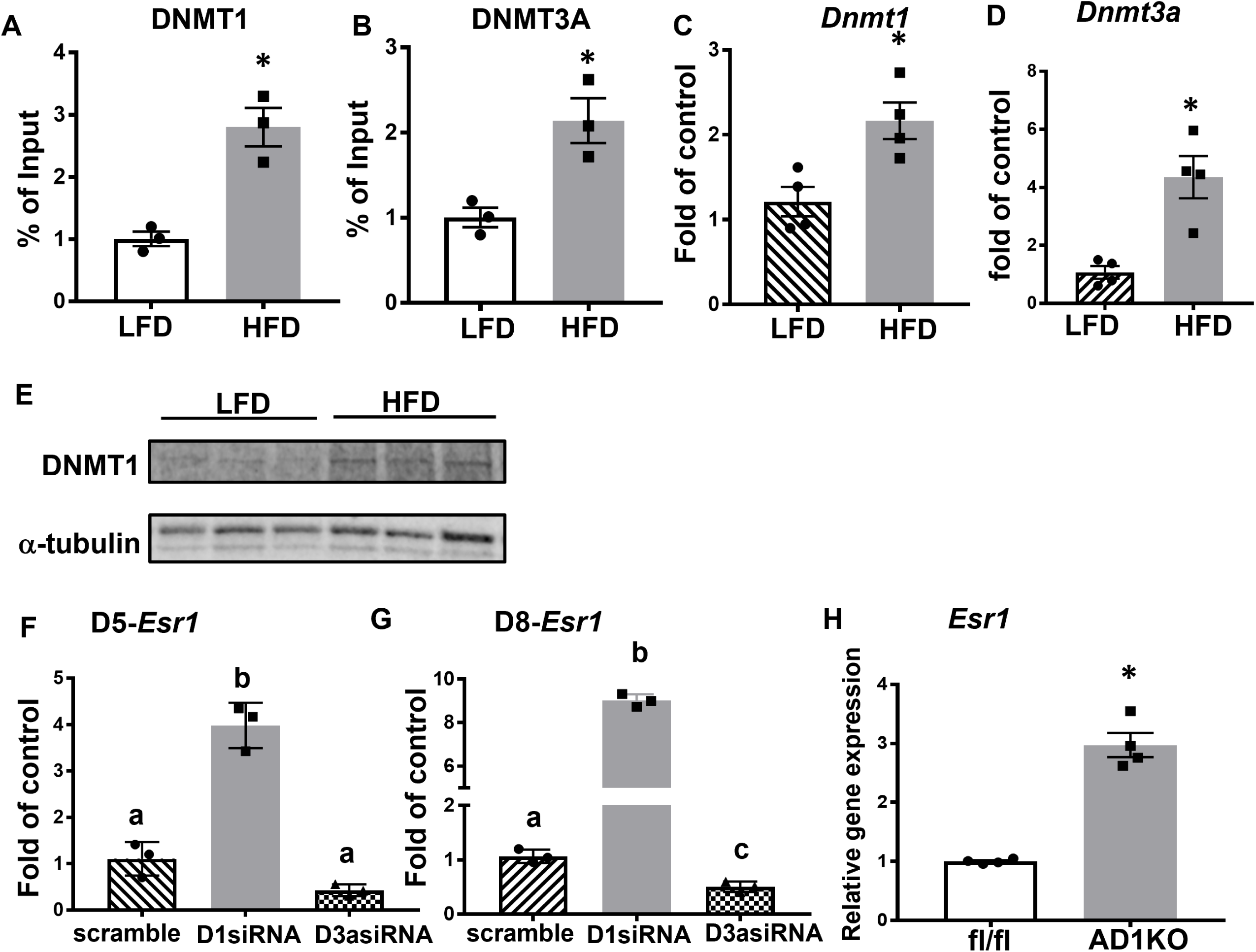
DNMT1 mediates HFD-induced increase of DNA methylation at *Esr1* promoter. (A-B) ChIP analysis of binding of DNMT1 (A) and DNMT3a (B) at *Esr1* promoter in gWAT of male mice fed with LFD or HFD for 12 weeks, n=3/group. *p<0.05 vs. LFD by Student’s t test. (C-D) Expression of *Dnmt1* (C) and *Dnmt3a* (D) in gWAT of male mice fed with LFD or HFD for 12 weeks, n=4/group. *p<0.05 vs. LFD by Student’s t test. (E) DNMT1 protein levels in gWAT of male mice fed with LFD or HFD for 12 weeks, n=3/group. (F-G) Expression of *Esr1* in 3T3-L1 adipocytes with *Dnmt1* or *Dnmt3a* knockdown at day 5 (F) or day 8 (G) of differentiation, n=3/group. Groups labeled with different letters are statistically different from each other as analyzed by ANOVA with Fisher’s LSD post hoc test. (H) Expression of *Esr1* in primary adipocytes differentiated from AD1KO and their fl/fl littermates, n=4/group. *p<0.05 vs. fl/fl by Student’s t test. All data are expressed as mean ± SEM.

### Inhibiting DNA methylation increases Esr1 expression and decreases inflammation in adipocytes

Since *Esr1* has been implicated in the regulation of inflammatory pathways (14-16), we interrogated whether inhibiting DNA methylation could increase *Esr1* expression and thereby decreasing inflammation in adipocytes. Indeed, knockdown of *Esr1* in 3T3-L1 adipocytes (**Suppl. Fig 4A**) promoted the expression of inflammatory genes such as tumor necrosis factor (*Tnfα)* and interleukin 1β (*Il1β)* (**Fig 3A**), while overexpression of *Esr1* (**Suppl. Fig 4B**) significantly suppressed these inflammatory genes expression (**Fig 3B**). We then investigated whether inhibiting DNA methylation pharmacologically by 5-aza-2’-deoxycytidine (5-aza-dC), a nucleoside-based DNA methyltransferase inhibitor that induces demethylation, would cause demethylation at the *Esr1* promoter and increase its expression, leading to an anti-inflammatory effect in adipose tissue. We fed 6-week-old male C57BL/6J mice with either LFD or HFD for 16 weeks to establish diet-induced obesity and then treated them with either saline or low dose 5-aza-dC (i.p) (0.25mg/kg BW, three times per week) for 6 weeks. This low dose 5-aza-dC treatment has been shown to exert very low toxicity as it did not change body weight and fat mass (10, 19). As expected, 5-aza-dC treatment did not change body weight and adiposity in these mice (**Suppl. Fig 5A-C**). However, while 5-aza-dC treatment did not change fed glucose levels, it significantly reduced fed insulin levels, and calculated glucose x insulin products (**Fig 3C**), indicating 5-aza-dC treatment improved insulin sensitivity in HFD-fed mice without changes in body weight and adiposity. This was confirmed by insulin tolerance test (ITT, **Fig 3C**). Interestingly, we found that HFD feeding suppressed the expression of *Esr1* in gWAT, which was restored by 5-aza-dC treatment (**Fig 3D)**. Moreover, 5-aza-dC treatment prevented HFD-induced expression of inflammatory genes such as *Tnfα* and *Il1β* (**Fig 3D**). This was further confirmed with a genetic approach using *Dnm1* deficient adipocytes differentiated from primary preadipocytes of AD1KO. Primarily differentiated adipocytes from *Dnmt1*-deficient or fl/fl mice were treated with saturated fatty acids and proinflammatory cytokines, two obesity-associated factors whose levels are commonly elevated in obesity. We found that the saturated fatty acids palmitate (C16) and stearate (C18) promoted inflammatory gene expression in adipocytes from fl/fl mice; this effect was prevented in adipocytes from AD1KO mice (**Fig 3E**). Similar results were observed in *Dnmt1*-deficient adipocytes treated with TNFα. *Dnmt1* deficiency significantly attenuated TNFα-induced expression of inflammatory genes, including *Tnfα*, interleukin 6 (*Il6*), inducible nitric oxide synthase 2 (*Nos2*/*iNos*), and monocyte chemoattractant protein 1 (*Mcp1*) (**Fig 3F**).

**Figure 3.**
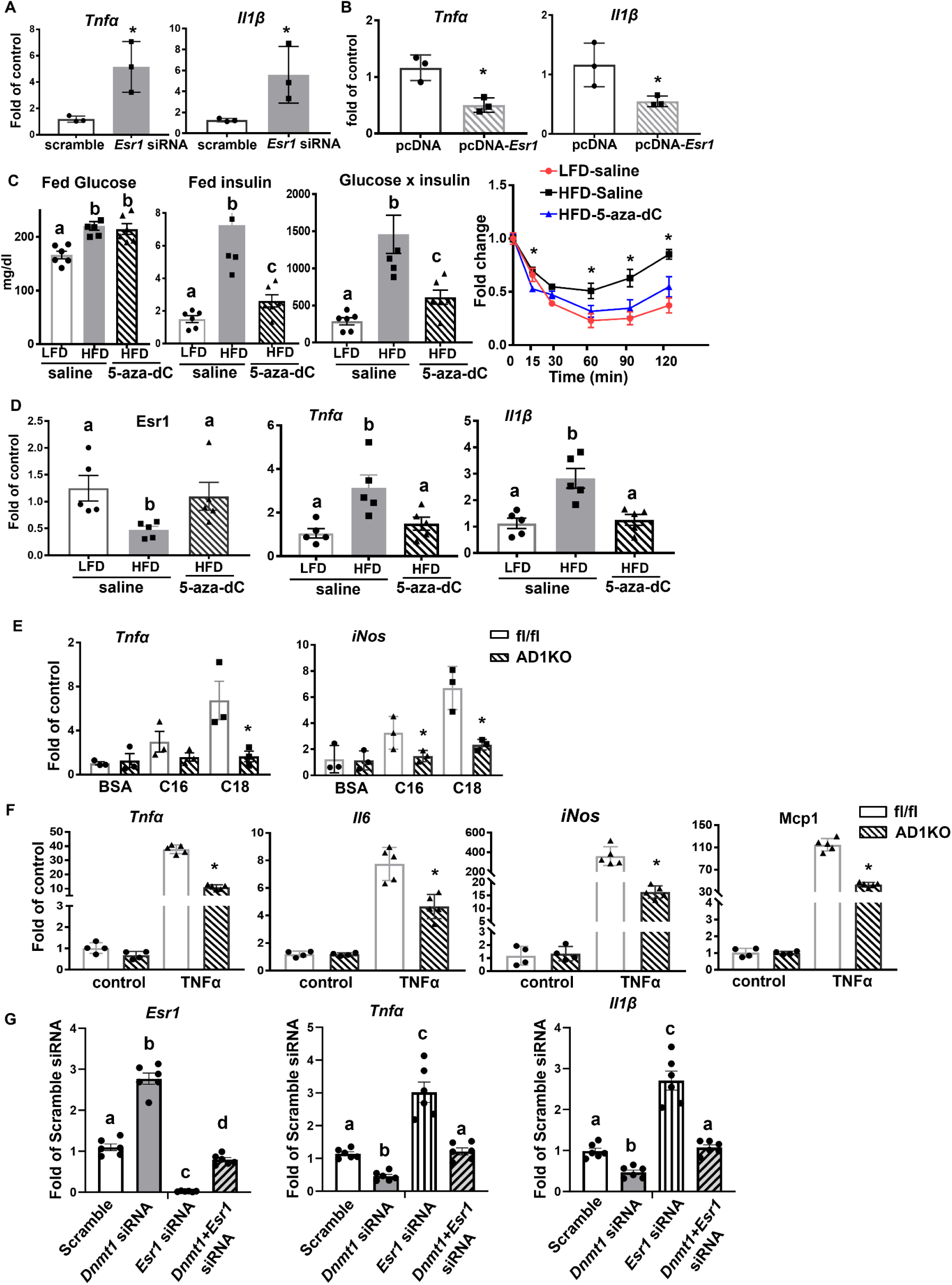
Inhibiting DNA methylation increases *Esr1* expression and decreases inflammation in adipocytes. (A-B) Pro-inflammatory gene expression in 3T3-L1 adipocytes with *Esr1* knockdown (A) or overexpression (B), n=3/group. *p<0.05 vs. Scramble in (A) or pcDNA in (B) by Student’s t test. (C) Plasma glucose, insulin, glucose x insulin products, and ITT in DIO mice treated with 5-aza-dC (0.25mg/kg BW, three times per week) for 6 weeks, n=4-6/group. Groups labeled with different letters are statistically different from each other; *p<0.05 vs. other groups. Statistical significance was analyzed by ANOVA with Fisher’s LSD post hoc test. (D) *Esr1*, and pro-inflammatory gene expression in gWAT of DIO mice treated with 5-aza-dC (0.25mg/kg BW, three times per week) for 6 weeks, n=5/group. Groups labeled with different letters are statistically different from each other as analyzed by Kruskal-Wallis non-parametric ANOVA by rank for *Esr1* and ANOVA with Fisher’s LSD post hoc test for *Tnfα* and *Il1β*. (E) Proinflammatory gene expression in adipocytes differentiated from AD1KO and fl/fl mice and treated with palmitate (C16) or stearate (C18), n=3/group. *p<0.05 vs. fl/fl by Student’s t test. (F) Proinflammatory gene expression in adipocytes differentiated from AD1KO and fl/fl mice and treated with TNFα, n=4-5/group. *p<0.05 vs. fl/fl by Student’s t test. (G) *Esr1*, *Tnfα* and *Il1β* expression in 3T3-L1 adipocytes with *Dnmt1* and/or *Esr1* knockdown. Day 5 differentiated 3T3-L1 cells were transfected with scramble, *Dnmt1*, *Esr1* or both *Dnmt1* and *Esr1* siRNA. Two days later, cells were treated with TNFα and samples were collected for gene expression analysis. n=6. Groups labeled with different letters are statistically different from each other as analyzed by ANOVA with Fisher’s LSD post hoc test. All data are expressed as mean ± SEM.

We have previously shown that inhibiting DNA methylation in macrophages by 5-aza-dC or *Dnmt1* deficiency exert significant anti-inflammatory effects (10). To further study whether the beneficial effect of inhibiting DNA methylation with either 5-aza-dC or adipocyte-specific *Dnmt1* deletion is mediated via adipocyte *Esr1*, we have knocked down both *Dnmt1* and *Esr1* in differentiated 3T3-L1 adipocytes. As shown in **Fig 3G**, knocking down *Dnmt1* significantly increased *Esr1* expression in 3T3-L1 adipocytes, with the concomitant downregulation of *Tnfα* and *Il1β*. As expected, *Esr1* knockdown in 3T3-L1 adipocytes significantly upregulated *Tnfα* and *Il1β* expression, and further abolished the inhibitory effects of *Dnmt1* knockdown on these proinflammatory cytokine expression, indicating that the beneficial effects of inhibiting DNA methylation on adipose tissue is mediated by adipocyte *Esr1*.

### Adipocyte-specific deletion of DNA methylation ameliorates HFD-induced obesity, adipose inflammation and insulin resistance

To more specifically study the role of adipocyte Dnmt1 in regulating adipose tissue inflammation, we have generated adipocyte-specific *Dnmt1* knockout (AD1KO) mice by crossing *Dnm1*-floxed mice with adiponectin-Cre mice. As expected, *Dnmt1* mRNA and protein levels were decreased in gWAT of AD1KO mice (**Suppl. Fig 6A-B**). We then further characterized the metabolic phenotypes of body weight, adipose inflammation and insulin sensitivity in both female and male AD1KO and their control fl/fl littermate mice fed HFD. **Fig 4A** showed that upon challenged with HFD, there was no significant difference in body weight in female mice between the two genotypes. However, female AD1KO mice exhibited a significant decrease in fat mass in various fat depots (**Fig 4B**) with smaller adipocytes (**Fig 4C**), whereas no difference in liver weight was observed (**Fig. 4B**). Using a PhenoMaster metabolic cage system, we found that female AD1KO mice exhibited higher energy expenditure (**Fig 4D**) with higher oxygen consumption (**Suppl. Fig 6C**) with no changes in locomotor activity and food intake (**Suppl. Fig 6D-E**), suggesting that the enhanced energy expenditure in female AD1KO mice largely accounted for their reduced adiposity. Further, female AD1KO mice also displayed improved glucose tolerance and insulin sensitivity as assessed by glucose and insulin tolerance tests (**Fig 4E** and **4F**). These data indicate that female mice with adipocyte Dnmt1 deficiency have reduced adiposity when fed HFD and are protected from obesity-induced insulin resistance.

**Figure 4.**
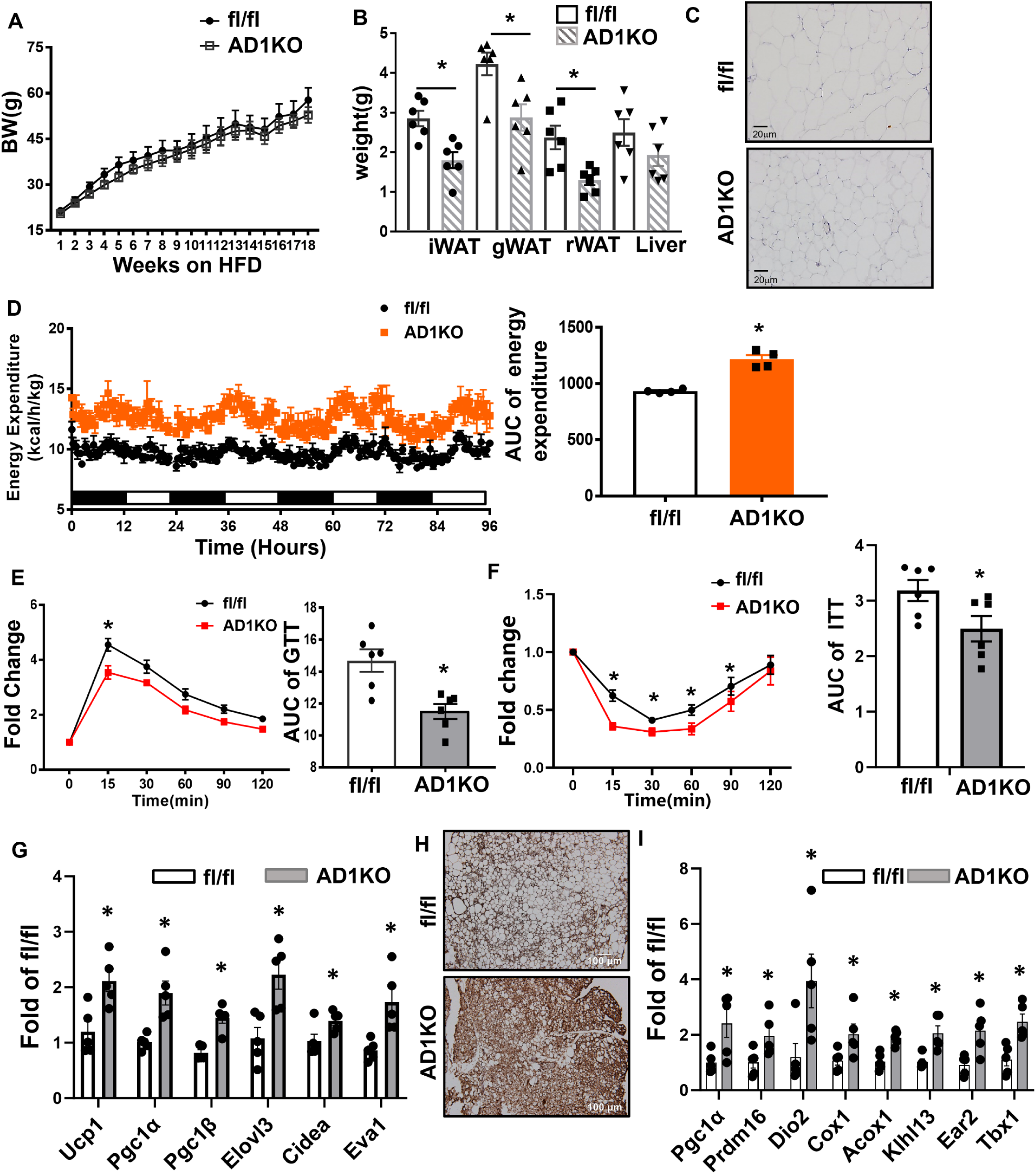
Adipocyte Dnmt1 deletion improves metabolic phenotypes in female mice fed HFD. (A-B) Body weight (A) and adipose tissue and liver weight (B) in female AD1KO and fl/fl mice fed HFD, n=6/group. *p<0.05 vs. fl/fl by Student’s t test. (C) H&E staining of gWAT of female AD1KO and fl/fl mice fed HFD. (D) Energy expenditure in female AD1KO and fl/fl mice fed HFD, n=4/group. *p<0.05 vs. fl/fl by Student’s t test. (E-F) GTT (E) and ITT (F) in female AD1KO and fl/fl mice fed HFD, n=6/group. *p<0.05 vs. fl/fl by ANOVA with repeated measures followed by Fisher’s LSD post hoc test. (G) Thermogenic expression in iBAT of female AD1KO and fl/fl mice fed HFD. n=5. *p<0.05 vs. fl/fl by Student’s t test. (H) UCP1 immunostaining in iBAT of female AD1KO and fl/f mice on HFD. (I) Thermogenic expression in gWAT of female AD1KO and fl/fl mice fed HFD. n=5. *p<0.05 vs. fl/fl by Student’s t test. All data are expressed as mean ± SEM.

To further study whether adipose tissue thermogenesis contributes to increased energy expenditure in female AD1KO mice, we have measured thermogenic gene expression in brown and white adipose tissues of female AD1KO and fl/fl mice. As shown in **Fig 4G**, the expression of thermogenic genes, including uncoupling protein 1 (*Ucp1*), peroxisome proliferative activated receptor γ coactivator 1 α (*Pgc1α*), *Pgc1β*, ELOVL fatty acid elongase 3 (*Elovl3*), cell death-inducing DNA fragmentation factor, α subunit-like effector A (*Cidea*) and epithelial V-like antigen 1 (*Eva1*) were upregulated in interscapular brown adipose tissue (iBAT). In consistence, immunohistochemistry staining with UCP1 antibodies also demonstrated increased UCP1 levels in iBAT of AD1KO mice (**Fig 4H**). In addition, the expression of several thermogenic genes and beige adipocyte marker, including *Pgc1α*, PR domain containing 16 (*Prdm16*), type 2 deiodinase (*Dio2*), cytochrome c oxidase subunit I (*Cox1*), acyl-Coenzyme A oxidase 1 (*Acox1*), kelch-like 13 (*Klhl13*), eosinophil-associated, ribonuclease A family, member 2 (*Ear2*) and T-box 1 (*Tbx1*), were also upregulated in gWAT of AD1KO mice (**Fig 4I**). These data suggest that adipocyte *Dnmt1* deletion promotes brown and beige adipocyte thermogenesis, which in turn contributes to increased energy expenditure observed in AD1KO mice.

Since *Dnmt1* deficiency suppresses adipocyte inflammation as shown above, we next examined adipose tissue inflammation in female AD1KO mice. *Dnmt1* deletion markedly decreased methylation rates on the individual CpG sites at the *Esr1* promoter as measured by pyrosequencing (**Fig 5A**). As expected, this was associated with an up-regulation of *Esr1* expression and a reciprocal down-regulation of inflammatory gene expression in gWAT of female AD1KO mice (**Fig 5B**). We further conducted RNA-seq analysis in gWAT in female AD1KO and their fl/fl littermate mice to thoroughly characterize inflammatory status through gene expression profiling. The volcano plot revealed a down-regulation of a panel of inflammatory/chemotactic genes in female *Dnmt1*-deficient gWAT (**Fig 5C**). This was consistent with the KEGG pathway analysis showing cytokine-cytokine receptor interaction, chemokine signaling pathway, and NF-kappa B signaling pathway among top ranked pathways identified (**Fig 5D**). Indeed, a hierarchical cluster analysis revealed a broad down-regulation of chemotactic (eg., chemokine (C-C motif) ligand 2 (*Ccl2/Mcp1*), Ccl3, Ccl4) and pro-inflammatory genes (e.g., *Tnfα*, *Il1β, Nos2/iNos*) in gWAT of female AD1KO mice on HFD (**Fig 5E**).

**Figure 5.**
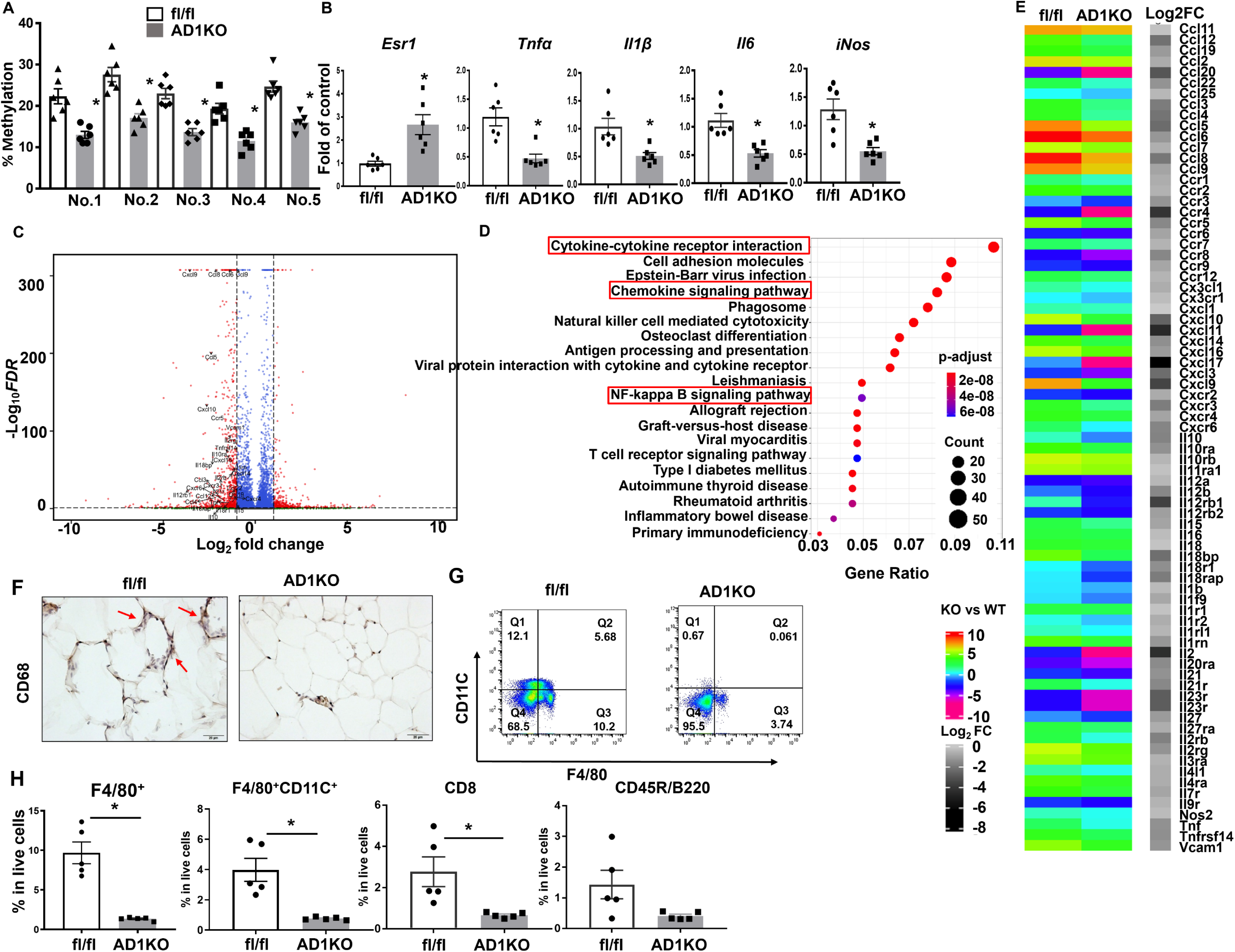
Adipocyte Dnmt1 deletion upregulates *Esr1* expression and reduces adipose tissue inflammation in female mice fed HFD. (A-B) *Esr1* promoter DNA methylation level (A) and mRNA expression level (B) in female AD1KO and fl/fl mice fed HFD, n=6/group. *p<0.05 vs. fl/fl by Student’s t test. (C-E) Volcano plot (C), pathway analysis (D) and heat map (E) analyzed from RNAseq data from gWAT of female AD1KO and fl/fl mice fed HFD. (F) CD68-immunostaining of gWAT of female AD1KO and fl/fl mice fed HFD. (G) FACS analysis of F4/80^+^CD11C^+^ ATMs in gWAT of female AD1KO and fl/fl mice fed HFD. (H) Percentage of F4/80^+^ ATMs, F4/80^+^CD11C^+^ ATMs, CD8^+^ T lymphocytes and CD45R/B220^+^ B lymphocytes in gWAT of female AD1KO and fl/fl mice fed HFD, n=5/group. *p<0.05 vs. fl/fl by Student’s t test. All data are expressed as mean ± SEM.

We further assessed the status of macrophage infiltration into adipose tissue in female AD1KO mice. Histochemical staining with antibodies against macrophage marker CD68 revealed a significant decrease of adipose tissue macrophage (ATM) content in gWAT of female AD1KO mice compared to that of fl/fl mice (**Fig 5F**). This was further confirmed by FACS analysis, which showed a significant reduction of F4/80^+^ ATMs, F4/80^+^/CD11C^+^ double positive ATMs and CD8^+^ T lymphocytes, as well as a tendency of decreased B220^+^ B lymphocytes in adipose tissue stromal vascular fraction cells in gWAT of female AD1KO mice compared to that of fl/fl mice (**Fig 5G-H**). These data suggest that adipocyte *Dnmt1* deletion reduces macrophage and lymphocyte infiltration in adipose tissue, thereby suppressing adipose inflammation.

We also characterized metabolic phenotypes of male AD1KO mice. There was no difference in body weight (**Suppl. Fig 7A**) and fat mass (**Suppl. Fig 7B**) between male AD1KO mice and their littermate control fl/fl mice. However, male AD1KO mice displayed a slight improvement in GTT test (**Suppl. Fig 7C-D**). Quantitative PCR analysis also showed a slight decrease in the expression of *Il6* (**Suppl. Fig 7E**).

We have also generated adipocyte-specific *Dnmt3a* knockout (AD3aKO) mice by crossing *Dnmt3a*-floxed mice (20) with adiponectin-Cre mice (18), and observed around 75% reduction of *Dnmt3a* expression in gWAT of AD3aKO mice (**Suppl. Fig 8A**). HFD-fed female AD3aKO mice did not show difference in body weight, fat pad weight, glucose tolerance/insulin sensitivity, *Esr1* expression, and inflammatory gene expression in WAT (**Suppl. Fig 8B-E**). There was also no difference in body weight between HFD-fed male AD3aKO and fl/fl mice (**Suppl. Fig 9A**). However, HFD-fed male AD3aKO mice had slightly improved glucose tolerance and insulin sensitivity (**Suppl. Fig 9B-C**) with no changes in the expression of *Esr1* and inflammatory genes in gWAT (**Suppl. Fig 9D**).

### Targeted methylation at the Esr1 promoter regulates adipocyte inflammation/chemotaxis and insulin sensitivity

Our data thus far suggest a key role of DNMT1 in the regulation of adipocyte inflammation and insulin sensitivity; however, it is not clear whether specific methylation at the *Esr1* promoter mediates the effect of DNMT1 in these processes. In addition, DNMT1 may have many downstream targets that confound the metabolic phenotypes observed in AD1KO mice. We therefore employed a modified CRISPR/RNA-guided system (21, 22) to specifically induce methylation/demethylation at the *Esr1* promoter. We first tested several single guide RNAs (sgRNAs) that surround the CpG sites at the *Esr1* promoter. 3T3L1 adipocytes were infected with lentivirus expressing sgRNAs (Scramble or *Esr1 targeting*) and dCas9-DNMT3a or dCas9-TET1 with *Esr1* mRNA as the readout. We identified one sgRNA (S7) that mediated the most potent inhibition or stimulation of *Esr1* expression for dCas9-DNMT3a and adCas9-TET1, respectively (**Suppl. Fig 10**).

To mimic the physiological milieu of adipose tissue where adipose tissue macrophages (ATMs) are recruited to fat tissue and constantly interact with adipocytes in a paracrine fashion, we examined adipocyte chemotaxis by a macrophage migration assay. We employed a co-culture system where RAW264.7 murine macrophages were grown in transwell inserts, which were then placed into lower well chambers containing differentiated L1 adipocytes infected with lentivirus expressing dCas9-DNMT3a or dCas9-TET1 along with S7 sgRNA. We found more macrophage migration in transwells cocultured with adipocytes infected with dCas9-DNMT3a and S7 sgRNA (**Fig 6A**). Meanwhile, the adipocytes infected with lentivirus expressing dCas9-DNMT3a and S7 sgRNA displayed enhanced proinflammatory gene expression than control adipocytes expressing scramble non-targeting sgRNA (**Fig 6B**). In contrast, we observed less macrophage migration in transwells cocultured with adipocytes infected with dCas9-TET1 and S7 sgRNA (**Fig 6C**). The adipocytes expressing dCas9-TET1 and S7sgRNA exhibited a significant decrease in proinflammatory gene expression (**Fig 6D**). These data suggest that DNA methylation at the *Esr1* promoter regulates adipocyte inflammation and chemotaxis.

**Figure 6.**
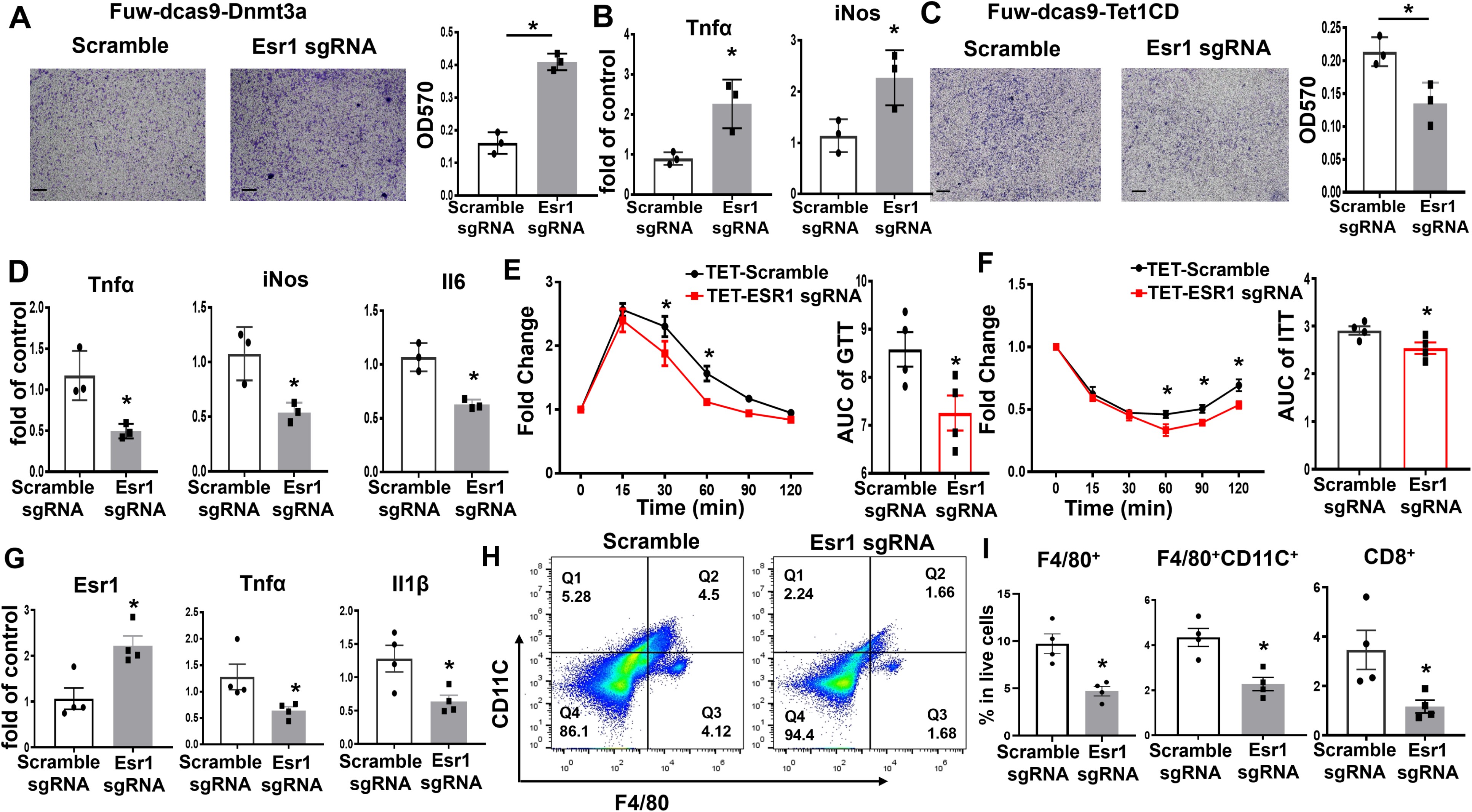
Targeted methylation at the *Esr1* promoter regulates adipocyte inflammation/chemotaxis and insulin sensitivity. (A-B) Macrophage migration assay (A) and inflammatory gene expression (B) in RAW264.7 macrophages co-cultured with 3T3-L1 adipocytes infected with lentivirus expressing scramble sgRNA or S7 sgRNA targeting *Esr1* promoter along with dCas9-DNMT3a, n=3/group, *p<0.05 vs. Scramble by Student’s t test. (C-D) Macrophage migration assay (C) and inflammatory gene expression (D) in RAW264.7 macrophages co-cultured with 3T3-L1 adipocytes infected with lentivirus expressing scramble sgRNA or S7 sgRNA targeting *Esr1* promoter along with dCas9-TET1, n=3/group, *p<0.05 vs. Scramble by Student’s t test. (E-F) GTT (E) and ITT (F) in HFD-fed female C57BL/6J mice surgically injected with dCas9-TET1 along with either scramble sgRNA or S7 sgRNA targeting *Esr1* promoter in gWAT, n=4. *p<0.05 vs. Scramble sgRNA by ANOVA with repeated measures followed by Fisher’s LSD post hoc test. (G-I) Gene expression (G), FACS analysis of F4/80^+^CD11C^+^ ATMs (H), and percentage of F4/80^+^ ATMs, F4/80^+^CD11C^+^ ATMs, and CD8^+^ T lymphocytes (I) in gWAT of HFD-fed female C57BL/6J mice surgically injected with dCas9-TET1 along with either scramble sgRNA or S7 sgRNA targeting *Esr1* promoter in gWAT, n=4. *p<0.05 vs. Scramble sgRNA by Student’s t test. All data are expressed as mean ± SEM.

To further investigate the role of DNA methylation at the *Esr1* promoter in the regulation of adipocyte inflammation and insulin sensitivity *in vivo*, we surgically injected lentivirus expressing dCas9-TET1 plus either the targeting S7 sgRNA or the scramble non-targeting sgRNA into gWAT of female C57/BL6J mice. Animals were then put on HFD for 12 weeks and were characterized for their metabolic phenotypes and adipose tissue inflammation. We successfully infected the fat depots with lentivirus expressing dCas9-TET1 as shown by mCherry IHC staining (**Suppl. Fig 11A**). Mice injected with lentivirus expressing dCas9-TET1 and S7 sgRNA had no difference in body weight (**Suppl. Fig 11B**); however, they exhibited improved glucose tolerance and insulin sensitivity as assessed by glucose and insulin tolerance tests compared to mice injected with lentivirus expressing dCas9-TET1 and scramble sgRNA (**Fig 6E** and **6F**). Moreover, lentiviral infection with dCas9-TET1 and S7 sgRNA increased *Esr1* gene expression in gWAT (**Fig 6G**). This was associated with decreased expression of inflammatory genes such as *Tnfα* and *Il1β* in gWAT (**Fig 6G**). FACS analysis also revealed a significant reduction of F4/80^+^ ATMs, F4/80^+^/CD11C^+^ double positive ATMs, and CD8^+^ T lymphocytes in gWAT of HFD-fed mice injected with lentivirus expressing dCas9-TET1 and S7 sgRNA (**Fig 6H-I**).

## DISCUSSION

In this study, we tested the hypothesis that epigenetic regulation at the *Esr1* promoter by HFD feeding, which often causes obesity, mediates adipocyte inflammation and chemotaxis, leading to obesity-induced insulin resistance and type 2 diabetes. The premise of this hypothesis stems from several prior observations. For one, chronic inflammation is a key link between obesity and insulin resistance/type 2 diabetes and adipose tissue plays an important role in obesity-induced inflammation and insulin resistance (1, 2). Second, inflammatory signaling pathways in adipose tissue can be activated in nutrient-rich conditions (e.g. HFD feeding) (3). However, it is not fully understood how HFD feeding alters the inflammatory program. Our study demonstrates that enhanced DNA methylation at the *Esr1* promoter by HFD plays an important role in mediating exaggerated adipose chemotaxis and inflammation, which may contribute to obesity-induced insulin resistance and type 2 diabetes.

Estrogen receptors (ESRs or ERs), members of nuclear receptor family, play a pivotal role in various metabolic and developmental processes, including metabolism, glucose homeostasis, differentiation, cell proliferation, immune-regulatory and anti-inflammatory function (14). There are two isoforms of ERs: ERα and ERβ, which are encoded by *Esr1* and *Esr2* genes, respectively (14). ERα (ESR1) is the predominant form in adipocytes (14). Evidence shows that ERα in the brain regulates food intake and energy metabolism (23). Mice with whole body deficiency of *Esr1* develop insulin resistance, suggesting a protective role of Esr1 in insulin sensitivity (24). In addition, *Esr1* has an anti-inflammatory function in immune cells due to its ability to interfere with the NFκB pathway (14). Recent studies have also demonstrated the anti-inflammatory role of *Esr1* in adipocytes (14-16). Genetic ablation of *Esr1* in adipocytes enhances adipose inflammation and macrophage infiltration, which might be mediated by increased *Mcp1* expression in adipocytes due to *Esr1* deficiency (15, 16). This may contribute to *Esr1*’s protective effect on insulin sensitivity (24). Here we discovered that HFD feeding significantly increased the binding of DNMT1 at the *Esr1* promoter, which resulted in increased DNA methylation at the *Esr1* promoter and subsequent downregulation of *Esr1* expression. Indeed, we found *Dnmt1* deficiency in adipocyte increased *Esr1* expression, resulting in decreased adipocyte chemotaxis. AD1KO mice with *Dnmt1* deficiency in adipocytes displayed increased *Esr1* expression, decreased adipose inflammation and improved insulin sensitivity upon HFD challenge. The anti-inflammatory effect of *Dnmt1* deficiency can be attributed to the decreased DNA methylation at the *Esr1* promoter, because specifically reducing DNA methylation at the *Esr1* promoter using a modified CRISPR/RNA-guided system increased *Esr1* expression, decreased adipose inflammation and improved insulin sensitivity in mice fed HFD, essentially recapitulating the phenotype of AD1KO mice.

Accumulating evidence suggests that epigenetic regulation plays a significant role in the development of obesity and its related disorders such as type 2 diabetes (25-29). However, relatively few studies address the role of epigenetics particularly DNA methylation in obesity-induced inflammation. We recently reported that inhibiting DNA methylation pharmacologically by 5-aza-2 deoxycytidine (5-aza-dC) ameliorated atherosclerosis in low density lipoprotein receptor knockout (*Ldlr*-/-) mice (30). This was associated with attenuated macrophage migration and adhesion to endothelial cells and reduced macrophage infiltration into atherosclerotic plaques (30). We also found that *Dnmt1* and *Dnmt3b* expressed in macrophages regulate macrophage polarization and inflammation, which further affects insulin sensitivity *in vivo* and *in vitro* (10, 31). Here we demonstrated a key role of adipocyte DNA methylation in mediating adipocyte chemotaxis and inflammation. Our data indicate that enhanced DNA methylation at the *Esr1* promoter by HFD suppresses *Esr1* expression in adipocytes, which may dampen its anti-chemotactic function and thus increase macrophage infiltration into adipose tissue. Meanwhile infiltrated macrophages in adipose tissue may undergo a phenotypic change toward to a more proinflammatory spectrum via an epigenetic mechanism involving DNA methylation. Enhanced DNA methylation at the *Pparγ* promoter by HFD decreases *Pparγ* expression in ATMs, promoting M1 macrophage polarization and inflammation (10, 31). It has been well documented that in the fat tissue, adipocytes and macrophages interact synergistically to generate inflammatory response and mediators in obesity (1, 32-35). Our study offers a novel understanding of adipose tissue inflammation from a unique perspective of epigenetic regulation.

Although DNMT1 is traditionally viewed as a maintenance enzyme that maintains DNA methylation patterns in dividing cells, several lines of evidence have pointed to a role for Dnmt1 beyond just maintaining methylation (7, 36). DNMT1 may also be involved in participation of *de novo* methylation on double-stranded DNA via a synergistic action along with other DNA methyltransferases (7, 37, 38). Since adiponectin-Cre we used to delete DNMT1 in adipocytes is primarily expressed in mature adipocytes (39), it is likely that DNMT1 exerts its pro-inflammatory function primarily by participating in *de novo* DNA methylation processes in adipocytes.

It is noteworthy that female AD1KO mice on HFD exhibited a stronger phenotype in improved insulin sensitivity and adipose inflammation than male AD1KO mice. Although the reason is not clear, estrogen as the ERα/ESR1 ligand may play a role. The presence of estrogen may synergistically activate the enhanced expression of *Esr1* caused by *Dnmt1* deficiency, leading to a more potent anti-inflammatory effect in adipocytes. Of note, while male AD1KO mice had little change in body weight, female AD1KO mice had a reduced adiposity upon HFD challenge, which may also contribute to their improved insulin sensitivity. The reduced adiposity in female AD1KO mice may be primarily due to their increased energy expenditure, as there was no change in food intake between HFD-challenged female AD1KO mice and fl/fl controls. Indeed, we found that HFD-challenged female AD1KO mice had significantly upregulated thermogenic markers in both brown and white adipose tissues, which could contribute to their increased energy expenditure. It is well documented that activation of ESR1 stimulates brown adipocyte thermogenesis and white adipocyte beiging (40, 41). Thus, our findings that increased *Esr1* expression in adipocytes due to *Dnmt1* deficiency could in turn promote brown and beige adipocyte thermogenesis in AD1KO mice, and contribute to their increased energy expenditure, reduced adiposity, and improved metabolic phenotypes.

Both brown and white adipose tissues are innervated by sympathetic nervous system that is important in regulating brown and beige adipocyte thermogenesis (42-45). Recent data suggest that adipose tissue derived neurotrophic factors and their receptors, including nerve growth factor (NGF) and its receptor tropomyosin receptor kinase A (TRKA)(46), brain-derived neurotrophic factor (BDNF) and its receptor tropomyosin receptor kinase B (TRKB) (47), and neurotrophic factor 3 (NT3) and its receptor tropomyosin receptor kinase C (TRKC) (48) are important in regulating SNS innervation in adipose tissue. Interestingly, it has been shown that estrogen and its receptor ESR1 promotes neuronal growth and development through interactions with the neurotrophins and their receptors (49), and estrogen stimulates the expression and production of these neurotrophic factors, including NGF, BDNF and NT3 in various brain regions and peripheral tissues (50-54). Thus, it is possible that promoting sympathetic nerve innervation in adipose tissue by estrogen and ESR1 via neurotrophic factor production may be another potential mechanism mediating ESR1’s effects on brown/beige adipocyte thermogenesis in AD1KO mice.

Along the course of our study, a recent report indicated that adipocyte DNMT3a mediates obesity-induced insulin resistance via increasing DNA methylation at the *Fgf21* gene (55). Taken together, DNA methylation catalyzed by DNMT1 and 3a may regulate adipocyte energy metabolism and inflammation via various pathways.

While our studies on the adipocyte DNMT1 are still ongoing, a recent study by Park et al showed opposite metabolic phenotype using a similar adipocyte-specific DNMT1 deficient mouse line (56). In the study by Park et al., adipocyte-specific DNMT1 deficient mice gained significant more weight even on a normal chow diet, and were further prone to diet-induced obesity when fed a high fat diet (56); whereas in our hands, our AD1KO mice had lower adiposity when fed a high fat diet, albeit with no changes in body weight. Although the exact reason for this discrepancy is not clear, several differences between the two studies may contribute to the different metabolic phenotypes observed. First, in the study by Park et al., metabolic studies were performed from only male mice, whereas in our study, we used both male and female AD1KO and fl/fl control mice. We did not find much metabolic difference between male AD1KO and fl/fl mice, thus, our study has primarily focused on female AD1KO mice. Second, there are several DNMT1-floxed mouse lines as well as adiponectin-Cre lines available according to Mouse Genome Informatics (MGI) (https://www.informatics.jax.org/allele/summary?markerId=MGI:94912, https://www.informatics.jax.org/allele/summary?markerId=MGI:106675). We obtained DNMT1-floxed mice from Mutant Mouse Regional Resource Centers (MMRRC) (*Dnmt1*-floxed line: MMRRC No. 014114)(17), which was created by inserting two loxP sites flanking exons 4 and 5, causing a frame shift and lacking the motifs for the catalytic domain. For the adiponectin-Cre line, we obtained from Jackson Laboratories (Stock #010803)(18), in which a BAC transgene was used to express Cre-recombinase under the control of adiponectin promoter. However, there was no information from Park et al regarding either the DNMT1-floxed mouse line or adiponectin-Cre line used in the study. It is possible that different DNMT1-floxed mouse lines and/or adiponectin-Cre lines used in the two studies may contribute to the discrepancy in mouse metabolic phenotypes observed. Third, differences in the genetic background of animals used in the two studies, or differences in diet and animal housing facilities may also contribute to the differences in the metabolic phenotypes observed in the two studies. Thus, further study is warranted to clarify the role of adipocyte DNMT1 in the regulation of adipocyte metabolic function.

In summary, we discovered that HFD feeding significantly enhanced DNA methylation at *Esr1* promoter in adipocytes, which is associated with down-regulation of *Esr1* expression. *Dnmt1* deletion in adipocytes increased *Esr1* expression, decreased adipose inflammation and improved insulin sensitivity in female mice with HFD challenge. Specifically reducing DNA methylation at the *Esr1* promoter using a CRISPR/RNA-guided system can largely recapitulate the phenotypes of adipocyte *Dnmt1*-deficient mice, suggesting a key role of DNA methylation at the *Esr1* promoter in regulating adipocyte chemotaxis and adipose inflammation. Deregulated epigenetic regulation, presumably by HFD, may result in altered adipocyte inflammation and chemotaxis, thereby contributing to obesity-induced inflammation and insulin resistance.

## MATERIALS AND METHODS

### Mice

Mice with adipocyte-specific DNA methyltransferase 1 (*Dnmt1*) or 3a (*Dnmt3a*) knockout (AD1KO or D3aKO) mice were generated by crossing *Dnmt1-* or *3a*-floxed mice (obtained from Mutant Mouse Regional Resource Centers (MMRRC); *Dnmt1*-floxed line: MMRRC No. 014114; *Dnmt3a*-floxed line: MMRRC No. 029885) with Adiponectin-Cre mice (Jackson Lab, Stock No. 010803)(18). The *Dnmt1*-floxed mouse was created by inserting two loxP sites flanking exons 4 and 5, which causes frame shift and lack the motifs for the catalytic domain (17). The *Dnmt3a*-floxed mouse was created by inserting two loxP sites flanking exon 19, which encodes the catalytic motif (20). For 5-aza-2’-deoxycytidine (5-aza-dC) treatment study, 6-week-old male C57BL/6J mice were fed either a low fat diet (LFD) (Research Diets D12450B, 10% calorie from fat) or a high fat diet (HFD) (Research Diets D12492, 60% calorie from fat) for 16 weeks to establish diet-induced obesity and were then randomly assigned to receive either saline or 5-aza-dC (0.25mg/kg) injection intraperitoneally (i.p) three times per week for up to 6 weeks. All the mice were housed in a temperature- and humidity-controlled animal research facility with a 12/12 h light–dark cycle and having *ad libitum* access to food and water. All animal studies were approved by the Institutional Animal Care and Use Committee at Georgia State University.

### Metabolic measurement

AD1KO, D3aKO mice and their respective floxed controls (fl/fl) were fed either LFD (Research Diets D12450B, 10% calorie from fat) or HFD (Research Diets D12492, 60% calorie from fat) diet for up to 20 weeks. The following metabolic measurements were conducted. **1)** Body weight was measured weekly and food intake of singly housed mice was measured over seven consecutive days. **2)** Blood glucose levels were measured by OneTouch Ultra Glucose meter (LifeScan, Milpitas, CA). Glucose tolerance and insulin tolerance tests (GTT and ITT, respectively) were conducted to determine glucose tolerance and insulin sensitivity as we previously described (10). **3)** Energy expenditure was measured using PhenoMaster metabolic cage systems (TSE Systems, Chesterfield, MO). At the end of HFD feeding studies, various white fat depots were dissected for further assessment of inflammatory status, including inflammatory gene expression, immunohistochemistry, and FACS analysis as described below.

### RNA extraction and quantitative RT-PCR

Total RNA from fat tissues or cultured adipocytes was isolated using the Tri Reagent kit (Molecular Research Center, Cincinnati, OH). The mRNA levels of the genes of interest were quantitated by a one-step quantitative RT-PCR with a TaqMan Universal PCR Master Mix kit (ThermoFisher Scientific, Waltham, MA) using an Applied Biosystems QuantStudio 3 real-time PCR system (ThermoFisher Scientific) as we previously described (10). The TaqMan primers/probes for all the genes measured were either purchased from Applied Biosystems (ThermoFisher Scientific) or commercially synthesized (Supplemental Table 1).

### Immunoblotting

Protein abundance in tissues was measured by immunoblotting as we described (10). Adipose tissue was homogenized in a modified radioimmunoprecipitation assay (RIPA) lysis buffer supplemented with 1% protease inhibitor mixture and 1% phosphatase inhibitor mixture (Sigma-Aldrich, St. Louis, MO). ysates were resolved by SDS-PAGE gels and were transferred to nitrocellulose membranes (Bio-Rad, Hercules, CA), which underwent blocking, washing, and sequential incubation with various primary antibodies and Alexa Fluor 680-conjugated secondary antibodies (Life Science Techenologies). The blots were developed with the Li-COR Imager System (Li-COR Biosciences, Lincoln, NE). The antibodies were listed in Supplemental Table 2.

### Immunohistochemistry (IHC)

Gonadal white adipose tissue (gWAT) was fixed in 10% neutral formalin, embedded in paraffin and sectioned at 5 µm thickness. The sections were immuno-stained with the primary and secondary antibodies, which were then developed with a peroxidase substrate from the DAB peroxidase substrate kit (Vector Labs, SK-4100). The primary and secondary antibodies are listed in Supplemental Table 2.

### Cell culture and SiRNA knockdown

3T3-L1 preadipocytes were cultured in a DMEM growth medium and were induced to differentiate into mature adipocytes with a differentiation cocktail as we previously described (57). Primary preadipocytes isolated from mouse gonadal white fat were cultured and differentiated as previously described (10). For SiRNA knockdown assays, day 5 or day 8 adipocytes were electroporated with targeting SiRNA or non-targeting scramble siRNA (GE Healthcare) using Amaxa Nucleofector II Electroporator (Lonza) with an Amaxa cell line nucleofector kit L (Lonza) as we previously described (58, 59), and cells were harvested 2-3 days after the transfection for further analysis.

### FACS analysis

FACS analysis of the macrophage content in adipose tissue was conducted as we previously described (10). Briefly, stromal vascular fraction (SVF) cells were isolated from epididymal fat using collagenase digestion, Cells were sequentially incubated with FcBlock (eBioscience) and APC-F4/80 (clone A3-1, AbD Serotec), PE-Cy7-CD11c (clone HL3, BD Pharmingen), APC-Cy7-CD8 (clone 53-6.7, BD Pharmingen), and Pacific blue-B220 (clone RA3-6B2, BD Pharmingen), followed by washing with the FACS buffer, fixing with paraformaldehyde and analyzing with BD FACSCalibur.

### Adipocyte chemotaxis

Adipocyte chemotaxis was assessed by a macrophage migration assay as we previously described (30, 60). Briefly, we employed an adipocyte-macrophage co-culture system where RAW264.7 macrophages were grown in transwell inserts, which were then placed into lower well chambers containing differentiated 3T3-L1 adipocytes with *Dnmt1* knockdown. Migrated cells were dissociated from the membrane, lysed, and quantified using CyQuant GR fluorescent dye with a Victor 3 plate reader (PerkinElmer).

### Esr1 promoter cloning and luciferase reporter assays

A 1kb fragment covering the *Esr1* proximal promoter and the 5’-untranslated regions enriched with CpG sites was PCR implied from mouse genomic DNA with primers listed in Supplemental Table 3. The PCR fragment was further subcloned into pGL3 basic luciferase reporter (Promega). The test to compare the activity of fully methylated vs. unmethylated Esr1 promoter was conducted as we previously described (10). The unmethylated promoter reporter was obtained by transforming the construct into *dam^–^/dcm^–^ E. coli* strain (New England Biolabs), while the fully methylated reporter was obtained by treating the construct with *SssI* methylase (New England Biolabs) in the presence of S-adenosylmethionine. The unmethylated or methylated *Esr1* reporter constructs were transfected into 3T3-L1 cells for luciferase activity measurement as we previously described (10).

### Reduced representation bisulfite sequencing (RRBS)

The whole genome DNA methylation was assessed using reduced representation bisulfite sequencing (RRBS) (11-13). Briefly, genomic DNA from the mouse epididymal white fat was extracted by the phenol-chloroform method and commercially sequenced by Beijing Genomics Institute (BGI) (Shenzhen, China). According to the instruction provided by the company, the genomic DNA was digested with the methylation-insensitive restriction enzyme *MspI*, ligated to sequencing adaptors, treated with sodium bisulfite, PCR-amplified for library construction and sequenced. The RRBS data analysis including Differentially Methylated Regions (DMRs), methylation rate and pathway analysis were conducted by BGI Bioinformatics Center or using the bioinformatics analysis pipeline as our co-author Dr. Huidong Shi described in (61). The methylation level at each CpG site was determined based on the number of sequences containing methylated CpGs versus the total number of sequences analyzed. For the comparison of DNA methylation rate differences between the HFD- and LFD-fed mice, data were summarized based on genomic features to generate tag density plots around transcription start and termination sites, exon-intron boundaries, CpG islands, and repeat elements and the data were uploaded to University of California Santa Cruz (UCSC) Genome Browser on Mouse (NCBI37/mm9) Assembly for methylated gene mapping as reported previously (61). Gene Ontology (GO) and Kyoto Encyclopedia of Genes and Genomes (KEGG) pathway analysis were performed with R package (v.3.2.0).

### RNA-seq analysis

Total RNA was isolated from gWAT as described above and was submitted to Beijing Genomics Institute (BGI, Shenzhen, China) for RNA-sequencing (RNAseq) analysis. Equal amount of RNAs from 4 animals/group were pooled and used for RNAseq analysis. Clean reads were aligned to the mouse reference genome (UCSC mm9). Differentially expressed genes between groups were defined as Log2 fold change ≥0.5 or ≤-0.5. Gene Ontology (GO) and Kyoto Encyclopedia of Genes and Genomes (KEGG) pathway analysis were performed with R package (v.3.2.0).

### Chromatin immunoprecipitation (ChIP) assays

ChIP assay was used to assess the binding of DNMTs to the *Esr1* promoter as we previously described (58). Fat tissue was fixed to isolate nuclei, which were sonicated to shear DNA. The DNA was immunoprecipitated with DNMT1 or DNMT3a antibodies, eluted, and followed by quantitative PCR using SYBR green. Primer sequences used in this study were shown in Supplemental Table 4.

### Bisulfite conversion and pyrosequencing

Pyrosequencing analysis was conducted to assess DNA methylation levels at *Esr1* promoter as we previously described (10). The genomic DNA from fat tissue was extracted by phenol/chloroform and followed by bisulfite conversion with an EpiTech Bisulfite Kit (Qiagen). The bisulfite-converted DNA was PCR-amplified, and the pyrosequencing was carried out by EpiGenDx (Hopkinton, MA). The pyrosequencing primers for the *Esr1* promoter are shown in Supplemental Table 5.

### Targeted DNA methylation at the Esr1 promoter

The mammalian lentiviral vectors FUW carrying dCas9-DNMT3a or dCas9-TET1, in which the catalytically inactive Cas9 (dCas9) has been engineered to be fused with the DNMT3a or TET1 (21, 22), were purchased from Addgene (Addgene No. 84476 and 84475). The guide RNA sequences targeting the CpG sites at the *Esr1* promoter was designed with GT-Scan website (http://gt-scan.braembl.org.au/gt-scan) and the targeting or non-targeting oligos were synthesized, annealed and cloned into the *AarI* sites of the pgRNA lentiviral vector (Addgene No. 44248). Lentivirus expressing dCas9-TET1-CD, dCas9-DNMT3a, or sgRNA was commercially produced by Vigene Biosciences, Inc. (Rockville, MD), and 10µl of lentivirus (1x10^5^TU/µl of titer) was surgically injected into mouse gonadal fat depots. Detailed primers information was listed in Supplemental Table 6.

### Statistical analysis

Student’s t-test, one-way or two-way analysis of variance (ANOVA) with Fisher’s Least Significant Difference (LSD) post-hoc test, or Kruskal-Wallis non-parametric ANOVA by rank were performed to evaluate statistical significance using GraphPad Prism version 5.0. In some experiments, one-way or two-way ANOVA with repeated measures was used for statistical analysis. Statistical significance was considered at p < 0.05. All data are shown as mean ± standard error (SEM).

## Author Contributions

RW performed experiments and data analysis; FL, SW, JJ, and XC assisted in experiments and data collection; HDS, XC, SW, YH, XZ, JAC, ZD and JS assisted in RRBS and RNAseq data analysis; LY contributed to study design, conceptual and technical inputs, and reviewed and edited the manuscript; BX and HS designed the study and wrote the manuscript.

## Acknowledgments

This work was supported in part by NIH grants R01DK118106 and R01DK125081 to B.X.; NIH grants R01DK115740, R01DK118106, R01DK116806 and R01DK130342 to H.S.; NIH grant R01DK116496, R01DK111052 and R01DK130342 to L.Y.

