## Supplemental Figures, Figure Legends and Tables for "Epigenetic Programming of Estrogen Receptor in Adipose Tissue by High Fat Diet Regulates Obesity-Induced Inflammation"

#### Supplemental Figure 1.

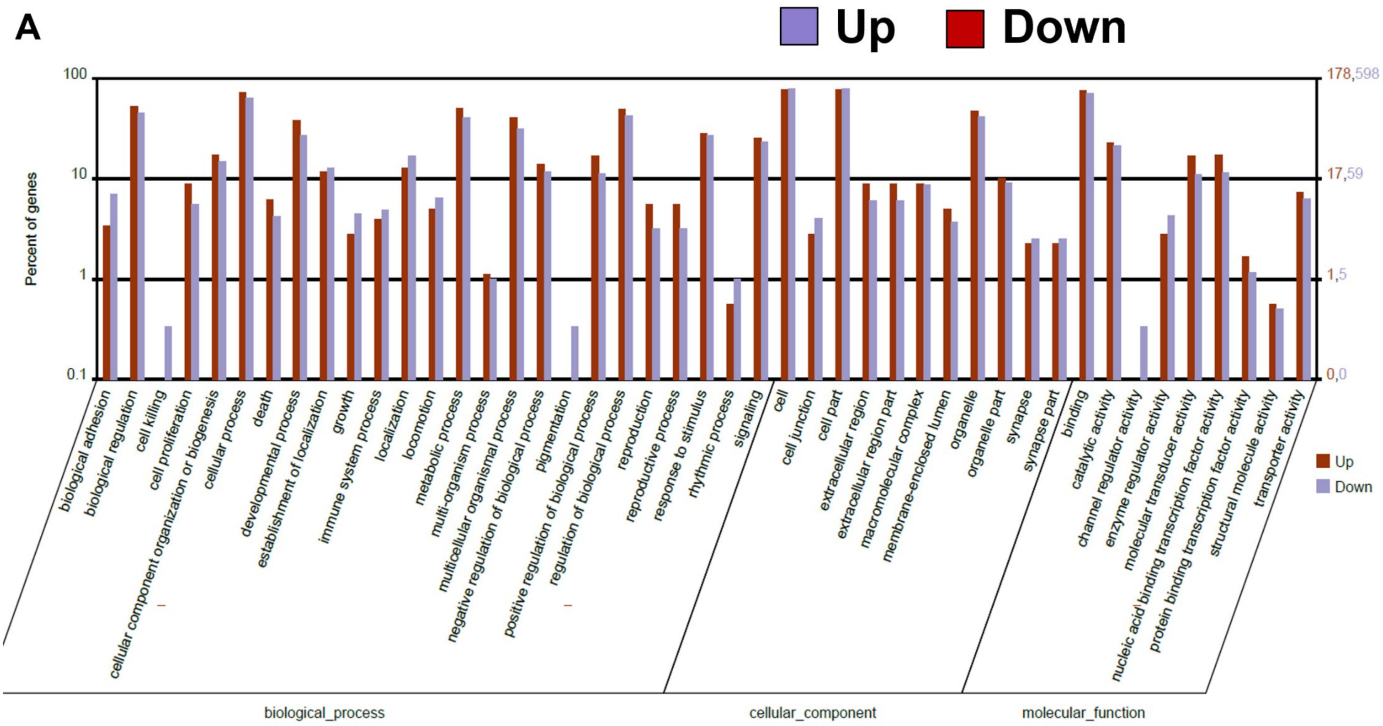


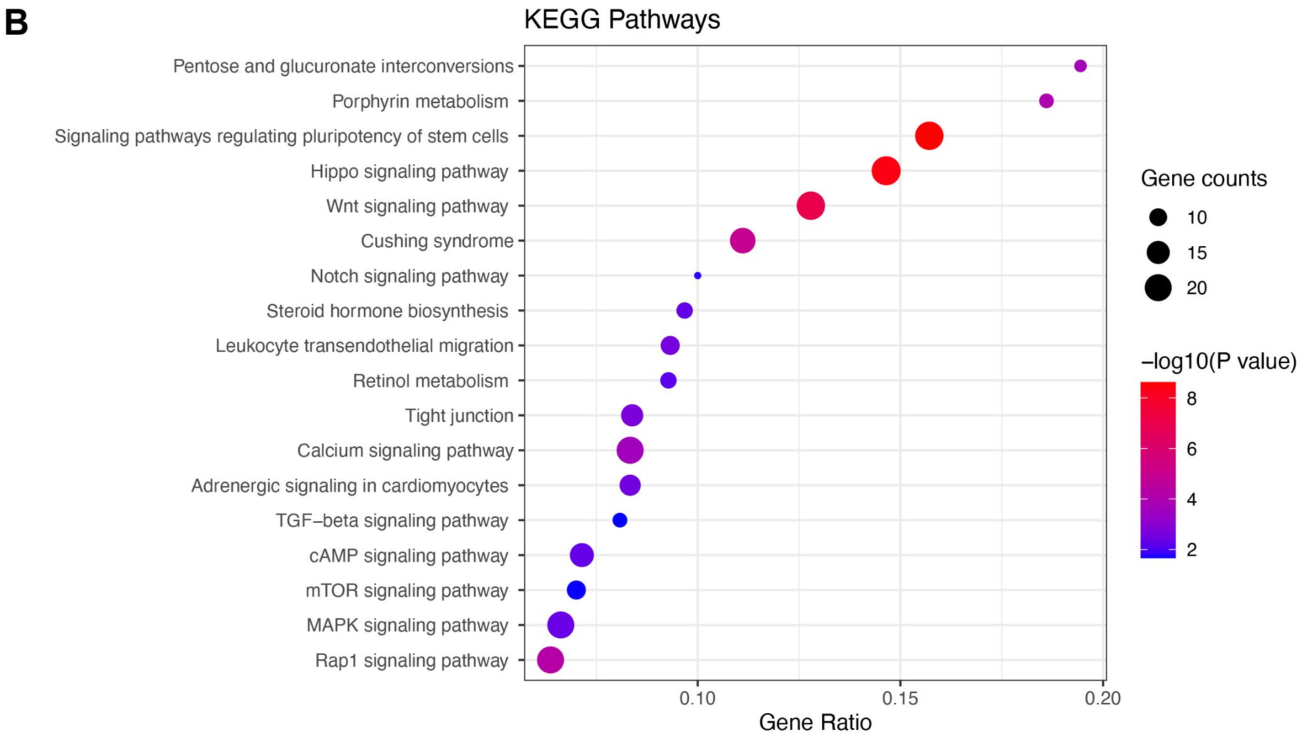


**Supplemental Figure 1.** (A-B) Gene Ontology (GO) analysis (A) and Kyoto Encyclopedia of Genes and Genomes (KEGG) analysis (B) of differentially methylated regions (DMRs) in gWAT of male mice fed with LFD or HFD for 12 weeks.

**Supplemental Figure 2.**

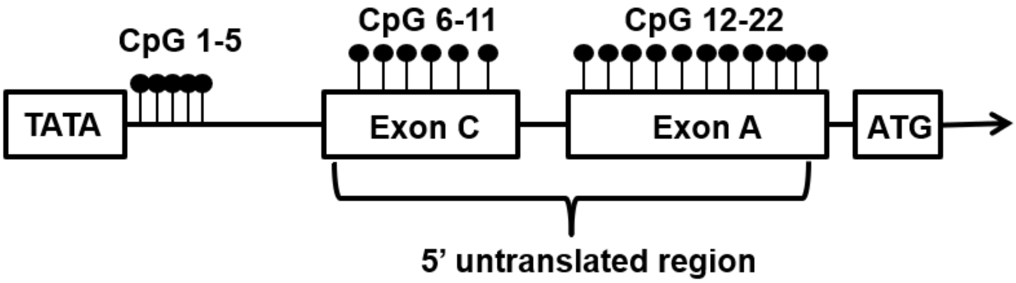


**Supplemental Figure 2.** Schematic illustration of the proximate promoter and 5'-UTR regions of *Esr1*. Closed circles indicate CpG sites on *Esr1* promoter.

**Supplemental Figure 3.**

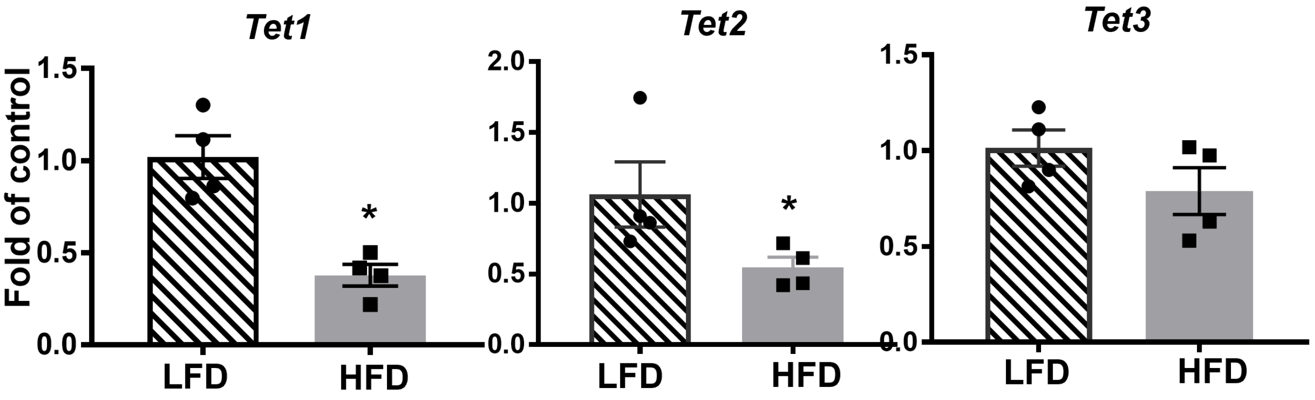


**Supplemental Figure 3.** Expression of *Tet1*, *Tet2* and *Tet3* in gWAT of male mice fed with LFD or HFD for 12 weeks. n=4/group. *$p < 0.05$ vs. LFD as analyzed by Student's t test. All data are expressed as mean ± SEM.

**Supplemental Figure 4.**

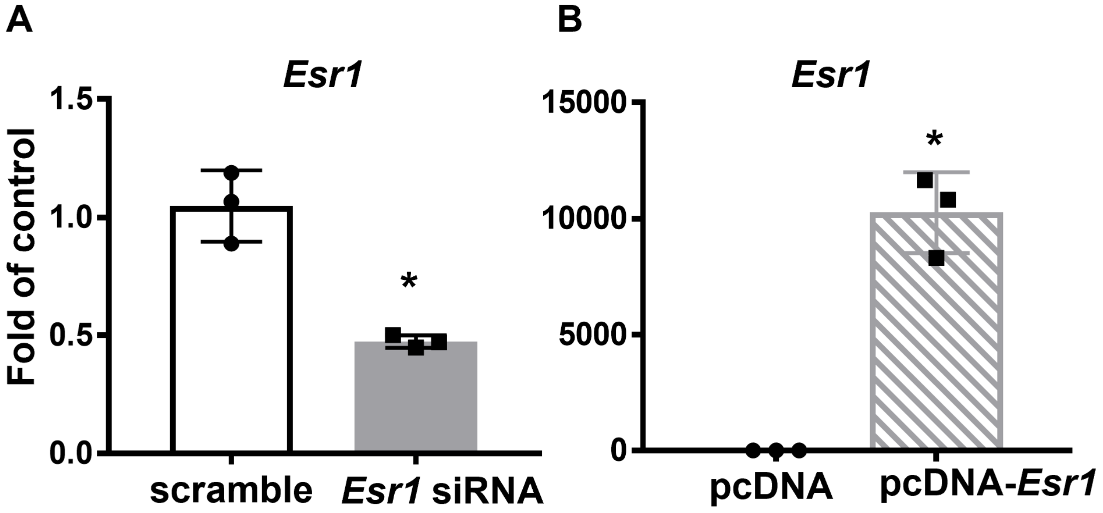


**Supplemental Figure 4.** *Esr1* knockdown (A) and overexpression (B) in 3T3-L1 adipocytes. n=3/group. *$p < 0.05$ as analyzed by Student's t test. All data are expressed as mean ± SEM.

#### Supplemental Figure 5.

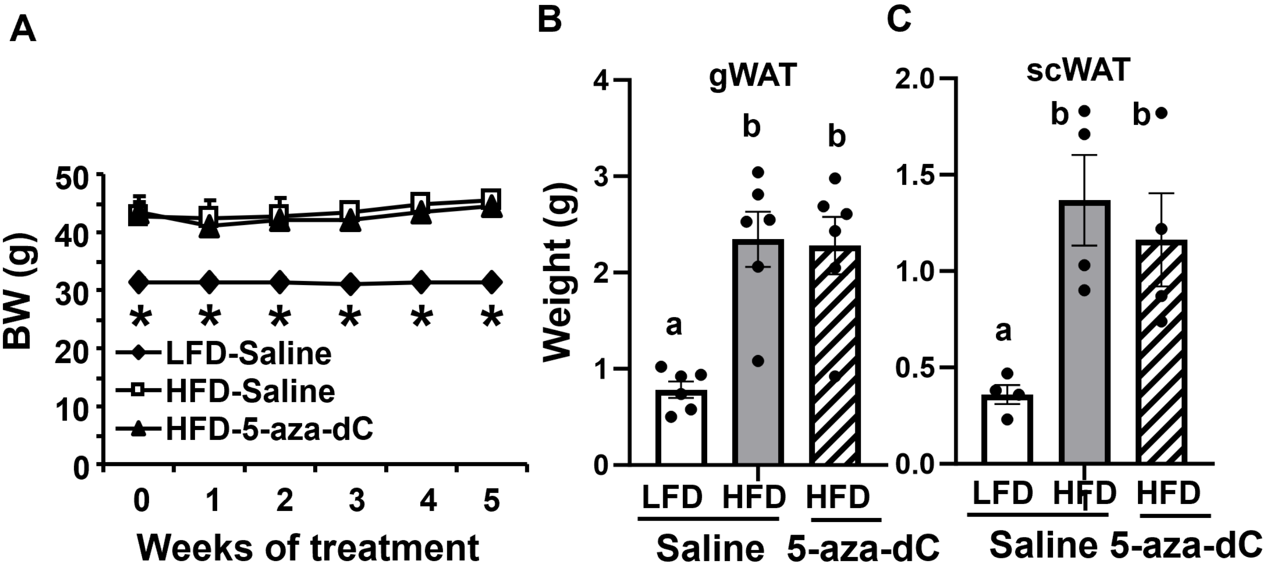


**Supplemental Figure 5.** 5-aza-dC treatment in DIO mice (0.25mg/kg BW, three times per week) for 6 weeks does not change body weight (A), gWAT (B) and scWAT (C) mass. n=4-6/group. In (A), *$p < 0.05$ vs. other groups. In (B), groups labeled with different letters are statistically different from each other. n=4-6/group. Statistical significance was analyzed by one-way ANOVA with Fisher's LSD post hoc test. All data are expressed as mean ± SEM.

#### Supplemental Figure 6.

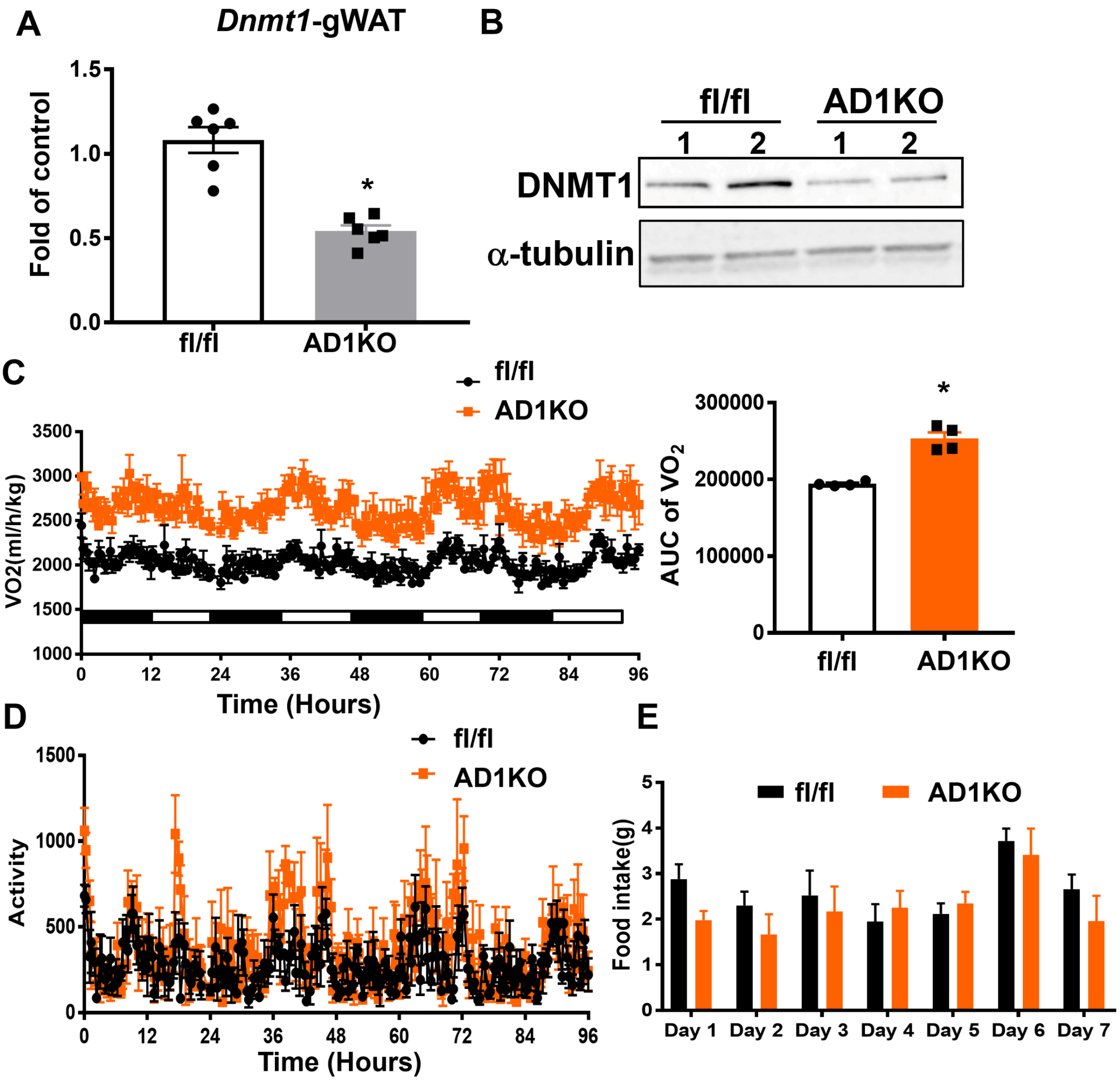

**Supplemental Figure 6.** Metabolic phenotypes in female AD1KO and their fl/fl littermate control mice fed HFD. (A-B) *Dnmt1* mRNA (A, n=6/group) and protein levels (B) in female AD1KO and fl/fl mice. *$p<0.05$ vs. fl/fl by Student's t test. (C-E) Oxygen consumption (C), locomotor activity (D) and food intake (E) in female AD1KO and fl/fl mice fed HFD, n=4/group. *$p<0.05$ vs. fl/fl by Student's t test. All data are expressed as mean ± SEM.

#### Supplemental Figure 7.

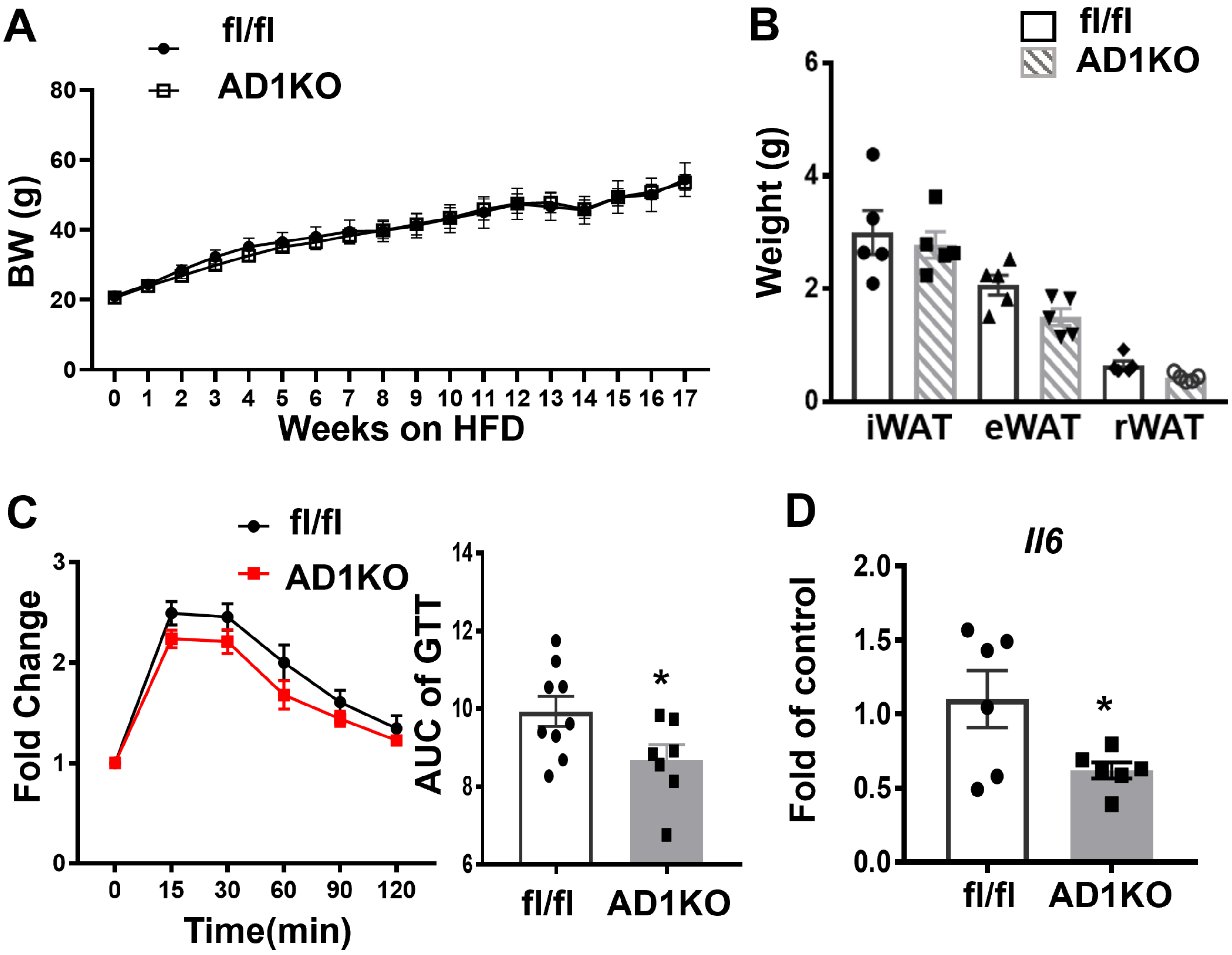


**Supplemental Figure 7.** Metabolic phenotypes in male AD1KO and their fl/fl littermate control mice fed HFD. (A-B) Body weight (A) and tissue weight of male AD1KO and fl/fl mice fed HFD, n=5/group. (C-D) GTT (C) and ITT (D) in male AD1KO and fl/fl mice fed HFD, n=7-9/group. (E) Proinflammatory gene expression in gWAT of male AD1KO and fl/fl mice fed HFD, n=5-6/group. *p<0.05 vs. fl/fl by Student's t test. All data are expressed as mean ± SEM.

### Supplemental Figure 8.

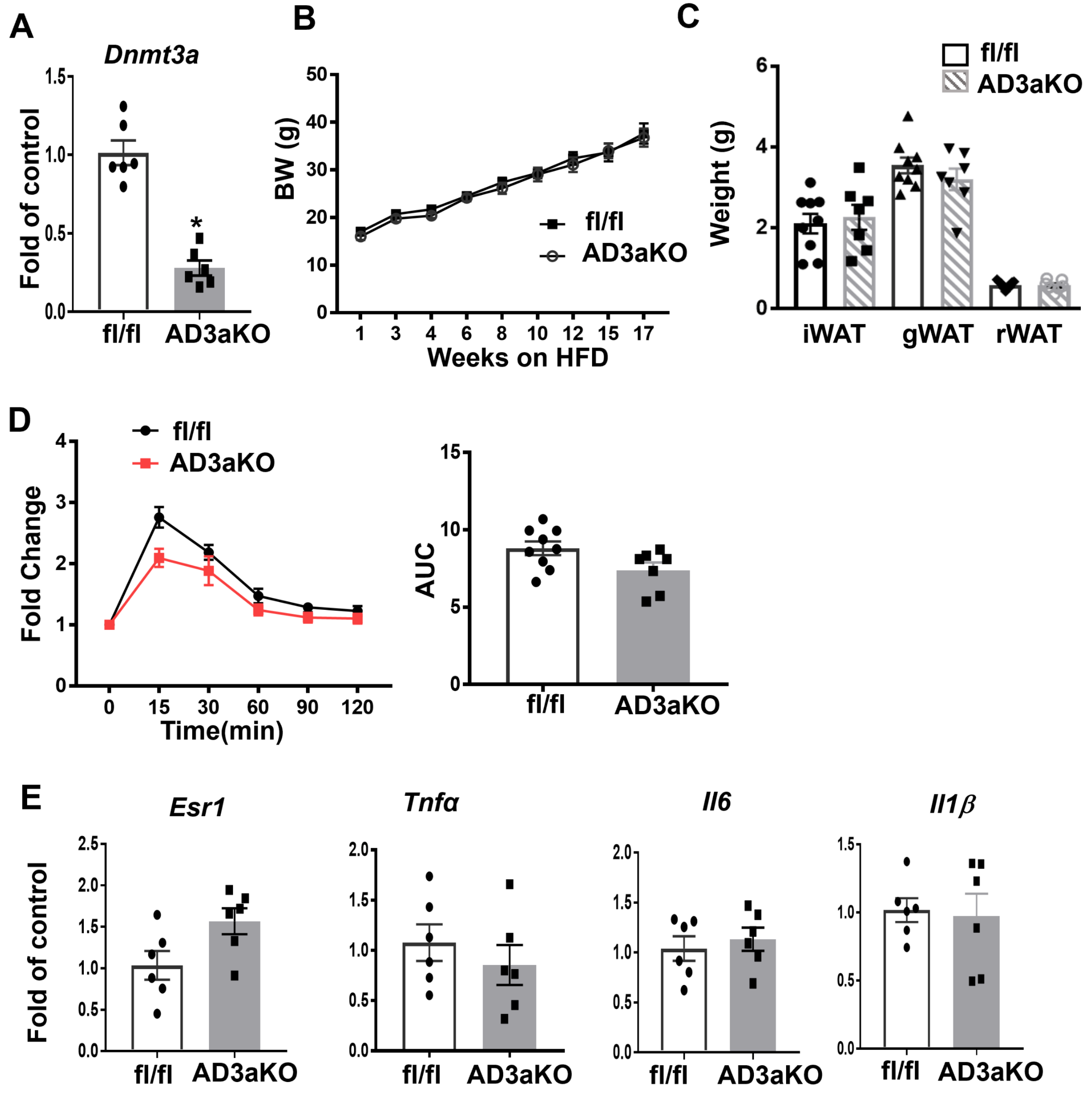

**Supplemental Figure 8.** Metabolic phenotypes in female AD3aKO and their fl/fl littermate control mice fed HFD. (A) *Dnmt3a* mRNA in female AD3aKO and fl/fl mice. n=6/group. *p<0.05 vs. fl/fl by Student's t test. (B-D) Body weight (B), tissue weight (C) and GTT (D) in female AD3aKO and fl/fl mice fed HFD, n=7-9/group. (E) Gene expression levels in gWAT of female AD3aKO and fl/fl mice fed HFD, n=6/group. All data are expressed as mean ± SEM.

### Supplemental Figure 9.

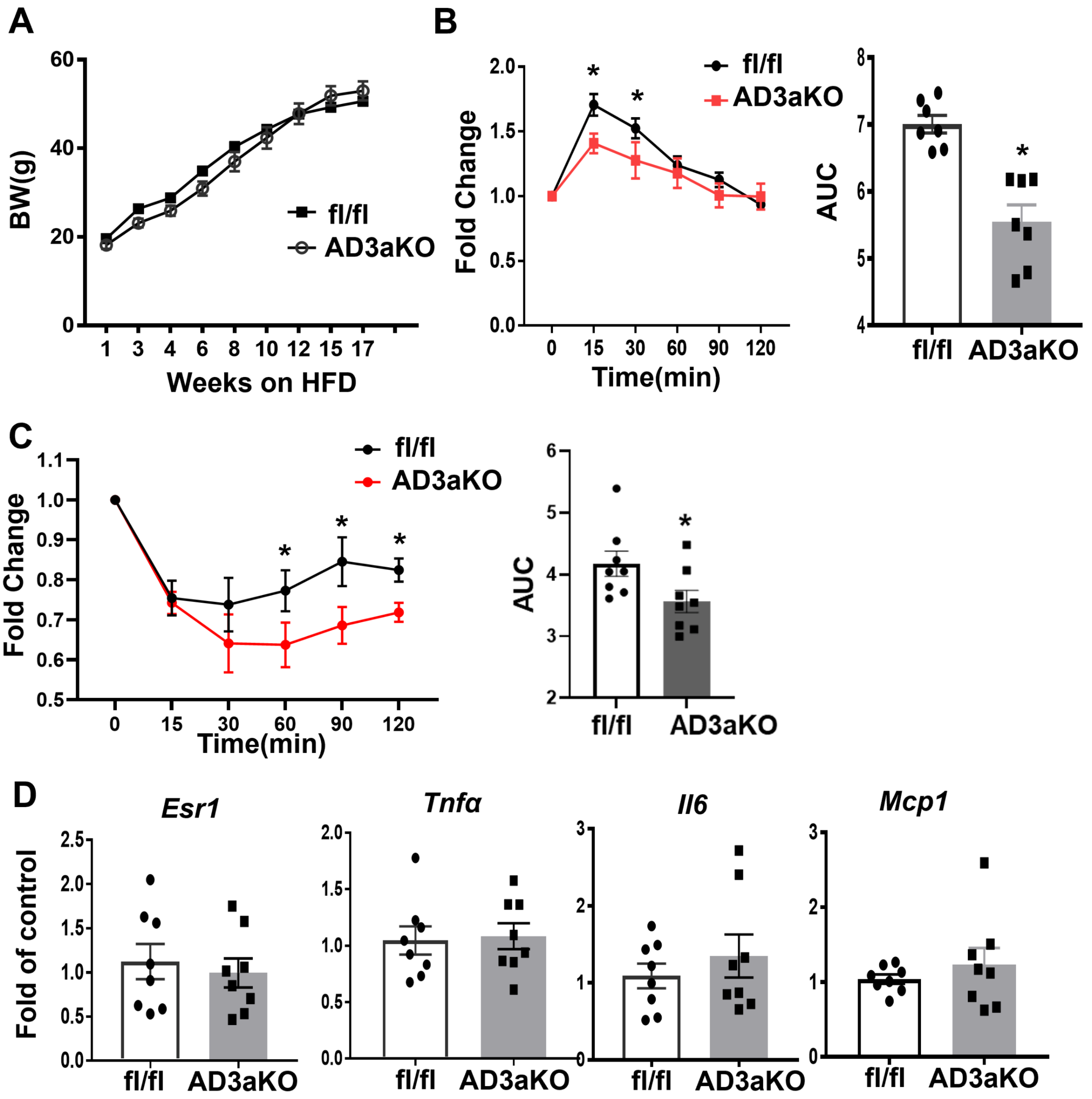

**Supplemental Figure 9.** Metabolic phenotypes in male AD3aKO and their fl/fl littermate control mice fed HFD. (A-C) Body weight (A), GTT (B) and ITT (C) in male AD3aKO and fl/fl mice fed HFD, n=7-8/group. *p<0.05 vs. fl/fl by Student's t test. (E) Gene expression levels in gWAT of female AD3aKO and fl/fl mice fed HFD, n=6/group. All data are expressed as mean ± SEM.

**Supplemental Figure 10.**

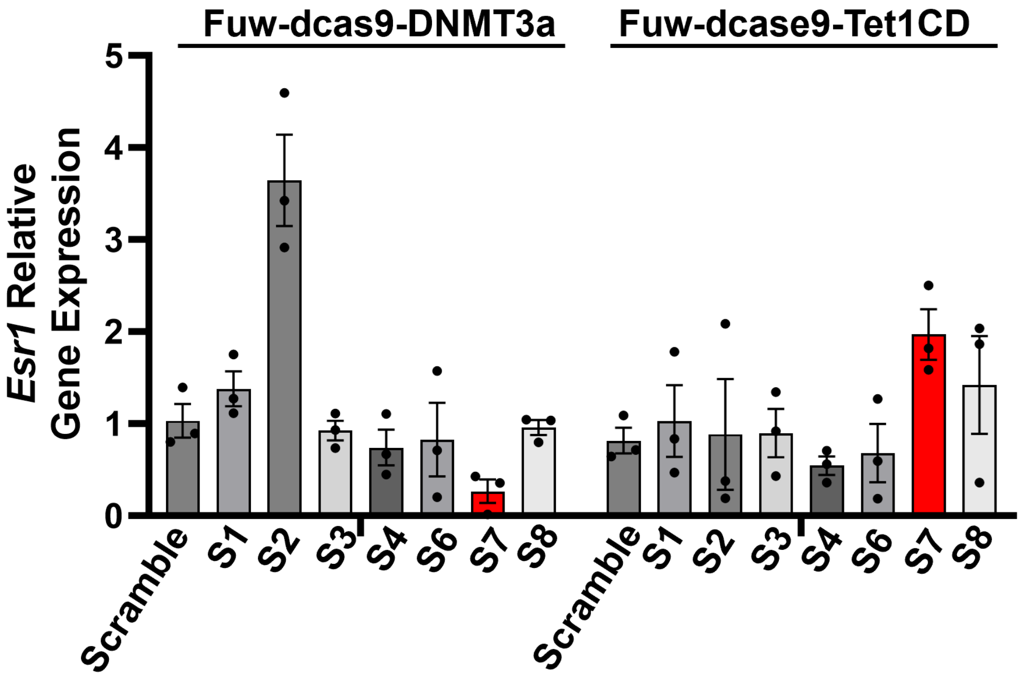


**Supplemental Figure 10.** *Esr1* expression in 3T3-L1 adipocytes infected with lentivirus encoding scramble sgRNA or sgRNAs targeting *Esr1* promoter together with either dCas9-DNMT3a or dCas9-TET1, n=3 *$p<0.05$ vs. Scramble by Student's t test. All data are expressed as mean ± SEM.

**Supplemental Figure 11.**

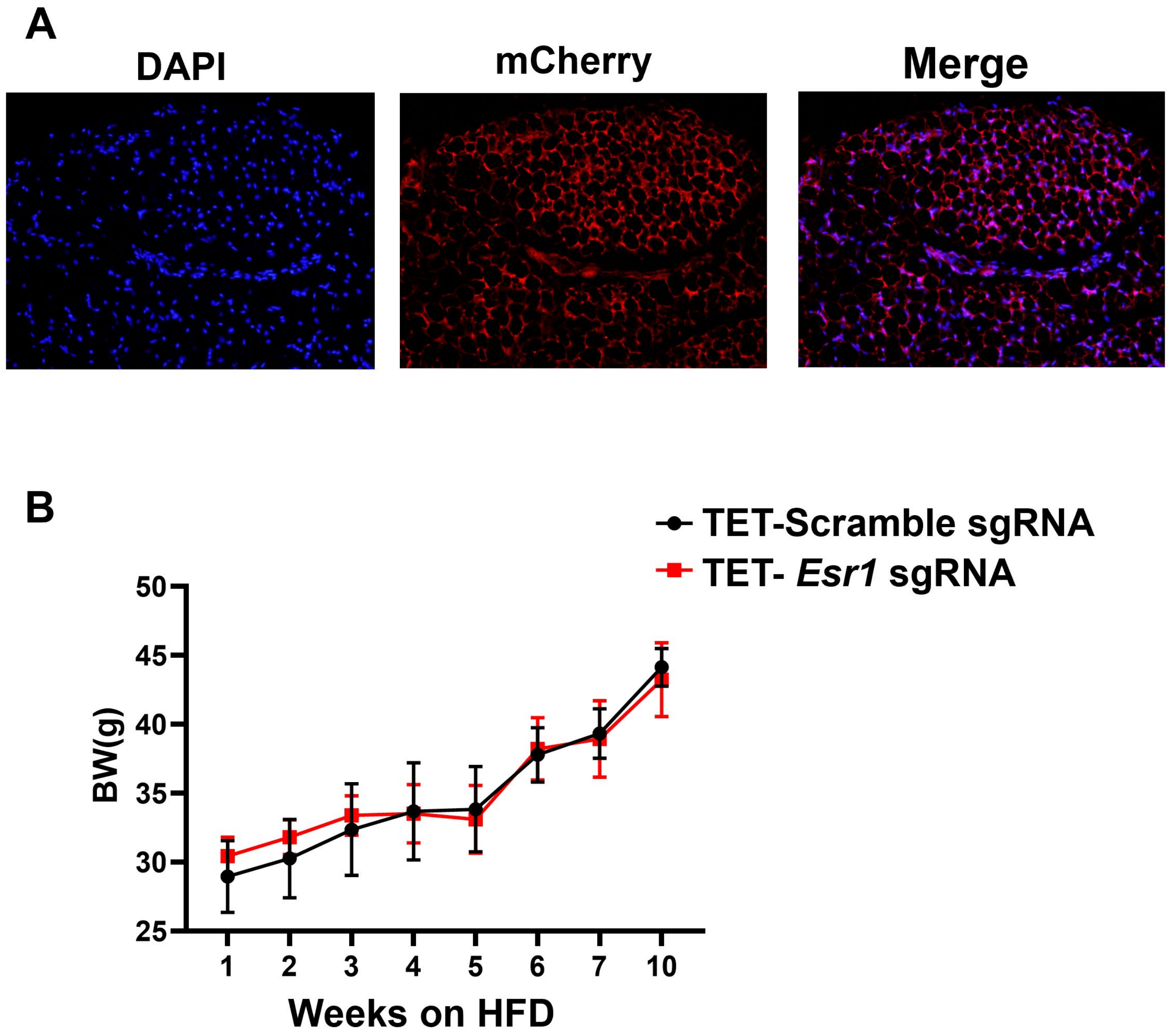


**Supplemental Figure 11.** (A) mCherry immunostaining in gWAT of female C57BL/6J mice surgically injected with dCas9-TET1 along with S7 sgRNA targeting Esr1 promoter into gWAT. (B) Body weight of HFD-fed female C57BL/6J mice surgically injected with dCas9-TET1 along with either scramble sgRNA or S7 sgRNA targeting Esr1 promoter, n=4. All data are expressed as mean ± SEM.

#### Supplemental Table 1. Primer/probe sets for gene expression

| Gene symbol | Company | Catalog # |
|---|---|---|
| *Esr1* | ABI | Mm00433149_m1 |
| *Tnf* | ABI | Mm00443258_m1 |
| *Nos2* | ABI | Mm00440502_m1 |
| *Il6* | ABI | Mm00446190_m1 |
| *Ccl2* | ABI | Mm00441242_m1 |
| *Il1β* | ABI | Mm01336189_m1 |
| *Dnmt1* | ABI | Mm00599783-g1 |
| *Dnmt3a* | ABI | Mm00432881_m1 |
| *Tet1* | ABI | Mm01169087_m1 |
| *Tet2* | ABI | Mm00524395_m1 |
| *Tet3* | ABI | Mm00805756_m1 |
| *Ucp1* | ABI | Mm01244861_m1 |
| *Pgc1α* | ABI | Mm01208835_m1 |
| *Pgc1β* | ABI | Mm00504730_m1 |
| *Elovl3* | ABI | Mm01194165_g1 |
| *Cidea* | ABI | Mm00432554_m1 |
| *Eva1* | ABI | Mm00468397_m1 |
| *Prdm16* | ABI | Mm00712556_m1 |
| *Dio2* | ABI | Mm0051664_m1 |
| *Cox1* | ABI | Mm04225243_g1 |
| *Acox1* | ABI | Mm01246834_m1 |
| *Klhl13* | ABI | Mm00470674_m1 |
| *Ear2* | ABI | Mm04207376_gH |
| *Tbx1* | ABI | Mm00448949_m1 |

#### Supplemental Table 2. Antibodies used in Immunoblotting, Immunohistochemistry and ChIP-qPCR

| Antibody | Company | Catalog# | Application |
|---|---|---|---|
| DNMT1 | Abcam | Ab87654 | WB |
| DNMT1 | Santa Cruz | sc-20701 | ChIP |
| DNMT3A | Santa Cruz | sc-373905 | ChIP |
| mCherry | Abcam | ab205402 | IF |
| CD68 | Abcam | Ab125212 | IHC |
| UCP1 | Abcam | Ab10983 | IHC |
| α-Tubulin | Santa Cruz | sc-53646 | WB |
| Goat anti-Mouse IgG (H+L) Highly Cross-Adsorbed Secondary Antibody, Alexa Fluor 680 | Invitrogen | A21058 | WB |
| Goat anti-Rabbit IgG (H+L) Highly Cross-Adsorbed Secondary Antibody, Alexa Fluor 680 | Invitrogen | A21109 | WB |
| Cy™3 AffiniPure Donkey Anti-Rabbit IgG (H+L) | Jackson ImmunoResearch | 711-165-152 | IF |

#### Supplemental Table 3. Primer sequences for *Esr1* promoter cloning

| Primer | Sequence (5'-3') |
|---|---|
| *Esr* promoter region-F1 | GGGGTACCCCAGACATTAAAACAACAACCCAGCA |
| *Esr* promoter region-R1 | GGAATTCC GCTAGCACCACCACTTGACT |
| *Esr* methylated region-F2 | GGAATTCCGGGCTGGAGTTTCTTCTAGGAAT |
| *Esr* methylated region-R2 | CCGCTCGAGCGGTGTATGTGGAGTGGCAGGGA |

#### Supplemental Table 4. Primer sequences for ChIP-qPCR assay

| Primer | Sequence (5’-3’) |
|---|---|
| *Esr*-F1 | GGGCTGGAGTTTCTTCTAGGAAT |
| *Esr*-R1 | CTGCACGAGCTGGATGAATG |

#### Supplemental Table 5. Primer sequences for pyrosequencing

| **Primer** | **Sequence (5'-3')** |
|---|---|
| *Esr1*-pyroseq-F1 | GGGGTTGGAGTTTTTTTTAGGAAT |
| *Esr1*-pyroseq-R1* | AACAACCAATAAACAAAACACTTAACA |
| *Esr1*-pyroseq-S1 | AGTTTTTTTTAGGAATGTTGA |
| *Esr1*-pyroseq-S2 | GTTTTTTAGTAGATAGTAAGGTT |

F: Forward primer;

R: reverse primer;

S: sequencing primer.

*Indicates primers with biotin-tag.

#### Supplemental Table 6. Primer sequences for guide RNA design

| Guide RNA | Primer (Forward: 5’-3’) | Primer (Reverse: 5’-3’) |
|---|---|---|
| Non-targeting gRNA | TTGGCCCCCGGGGGAAAAATTTTT | AAACAAAAATTTTTCCCCCGGGGG |
| *Esr1* gRNA | TTGGcgggaagcagccagtaggca | AAACtgcctactggctgcttcccg |
